# Serotonin GPCR-based biosensing modalities in yeast

**DOI:** 10.1101/2021.07.12.452006

**Authors:** Bettina Lengger, Emma E. Hoch-Schneider, Christina N. Jensen, Tadas Jakočiūnas, Emil D. Jensen, Michael K. Jensen

## Abstract

Serotonin is a key neurotransmitter involved in numerous physiological processes and serves as an important precursor for manufacturing bioactive indoleamines and alkaloids used in the treatment of human pathologies. In humans, serotonin sensing and signaling can occur by 12 G protein-coupled receptors (GPCRs) coupled to G proteins. To systematically assess serotonin GPCR signaling, we characterized reporter gene expression of a 144-sized library encoding all 12 human serotonin GPCRs in combination with 12 different Gα proteins in yeast exposed to serotonin. For the 5-HT4 receptor, we observe 25- and 64-fold changes in EC_50_ values and dynamic reporter gene outputs, respectively. Furthermore, we show that optimal biosensing designs enable high-resolution sensing of serotonin produced in yeast, as well as provide a platform for characterization of 19 serotonin GPCR polymorphisms found in human populations. Taken together, our study highlights serotonin biosensing modalities of relevance to both biotechnological and human health applications.

**Highlights:**

- Human serotonin G protein-coupled receptors display promiscuous Gα coupling in yeast
- Gα-coupled serotonin receptors display up to 64-fold changes in reporter expression output
- Differences in Gα protein evokes 25- and 2-fold difference in EC_50_ and sensitivity, respectively
- Serotonin receptor 5-HT4 and human SNP variants display physiologically relevant EC_50_ values in yeast
- 5-HT4 can be applied for high-resolution biosensing of serotonin produced from yeast

## Introduction

Serotonin is a monoamine neurotransmitter largely confined to the digestive and central nervous systems of humans, and implicated in a plethora of biological functions in humans, including mood, feelings, eating, and sleeping (Berger et al., 2009). In humans a total of 13 serotonin receptor genes and 1 pseudogene are found, and encodes for a total of 12 serotonin G protein-coupled receptors (GPCRs) and 1 ionotropic channel, together mediating serotonin signaling (Nichols and Nichols, 2008). Collectively, GPCRs are seven-transmembrane proteins, which allow cells to respond to extracellular stimuli by coupling the binding of a ligand to the activation of intracellular signaling pathways (Zhang et al., 2015). The intracellular signaling through GPCRs is mediated via a ligand-mediated conformational change serving as a guanine-exchange factor to activate heterotrimeric guanine nucleotide-binding protein (G protein), consisting of the three subunits Gα, Gβ and G□ (Syrovatkina et al., 2016). Binding of a ligand to the GPCR promotes a conformational change in the receptor, which in turn activates the GPCR-bound Gα subunit of the G protein. The exchange of Gα-bound GTP to GDP promotes the dissociation of the G protein from the GPCR as well as the separation of the Gα subunit from the Gβ□ dimer (Leberer et al., 1992). Following dissociation, the subunits relay intracellular signalling to ultimately effectuate an adequate transcriptional reprogramming in response to the extracellular milieu (Marinissen and Gutkind, 2001). While these modules constitute the core GPCR signalling, a dearth of knowledge challenges our understanding of how the approx. 800 GPCRs encoded in the human genome couple through 16 different Gα subunits (Pándy-Szekeres et al., 2018), notwithstanding the structure-affinity relationship between the great diversity of ligands and the GPCRs which have evolved to respond to them, including light, hormones, and small molecules, like serotonin ((Lengger and Jensen, 2020; UniProt Consortium, 2021)).

For more than three decades yeast has served as a platform for studying human GPCRs (Brown et al., 2000; King et al., 1990) with great potential in both medical and biotechnological application areas (Lengger and Jensen, 2020). The vast majority of GPCR studies in yeast are based on the mating pathway naturally activated by pheromone through the yeast mating-factor GPCRs Ste2/3 (Versele et al., 2001). Upon ligand activation, successful coupling of a heterologous GPCR with the yeast mating pathway can occur through coupling to the yeast Gα protein GPA1, which subsequently activates the mitogen-activated protein (MAP) kinase cascade consisting of Ste5/Ste7/Ste11, ultimately resulting in the activation of 100s of pheromone responsive genes (Lengger and Jensen, 2020). While a few studies successfully proved coupling of human GPCRs to GPA1 (Ehrenworth et al., 2017; King et al., 1990; Mukherjee et al., 2015), a key contribution to potentiate coupling of heterologous GPCRs to the yeast mating pathway was the discovery of improved coupling when exchanging the C-terminal five amino acids in the G□ part of GPA1 with complementary 5-mer human G□ signatures, also referred to as chimeric Gα proteins (Brown et al., 2000; Conklin and Bourne, 1993; Nakamura et al., 2015; Shaw et al., 2019). Likewise, knockout of *SST2*, a negative regulator of GPA1, and *FAR1*, an inducer of cell cycle arrest during mating, have been key steps to increase heterologous GPCR signalling in yeast, as demonstrated previously (Dohlman et al., 1996; Price et al., 1995), ultimately enabling the development of whole-cell biosensors based on >50 GPCRs (Kapolka et al., 2020; Lengger and Jensen, 2020; Shaw et al., 2019).

For serotonin GPCRs, three out of the twelve human serotonin GPCRs have successfully coupled to the yeast mating pathway, namely 5-HT1A (Brown et al., 2000), 5-HT1D (Brown et al., 2000; Nakamura et al., 2015) and the 5-HT4 receptors (Bean et al., 2021; Ehrenworth et al., 2017; Kapolka et al., 2020; Shaw et al., 2019). Likewise, in yeast, 10 mutants of the 5-HT1A receptor have been engineered to elucidate polymorphisms impacting receptor activation (Nakamura et al., 2015), while Kapolka *et al*. recently reported coupling of 5-HT4 with all 10 chimeric Gα proteins variants (Kapolka et al., 2020). Importantly, while yeast only provides a minimal platform for studying heterologous GPCR signalling through its mating pathway, physiologically relevant pharmacological properties as well as receptor specificity for the Gα chimera have shown to be consistent with EC_50_ values and cognate mammalian Gα protein coupling, respectively (Brown et al., 2000; Nakamura et al., 2015). Likewise, for biotechnological purposes, microbial production in yeast of a range of GPCR agonists with clinical applications can offer a solution for supply chain stability and scalability of production (Galanie et al., 2015; Luo et al., 2019). However, optimizing heterologous biosynthetic pathways for human bioactives in yeast using metabolic engineering is often a tedious endeavor, involving complex engineering to create optimal pathway designs for fermentation-based manufacturing of such bioactives. Here, GPCR-based serotonin biosensors have shown promising results with a 5-HT1A-based sensor coupled to the GPA1/Gαi3 chimera resulting in a sensor with 300% increase over basal fluorescence after activation with serotonin (Nakamura et al., 2015). Using a 5-HT4 based sensor, Ehrenworth et al. recently demonstrated that yeast-produced serotonin could be detected with a 2-fold change (Ehrenworth et al., 2017). Still, while serotonin receptors have been characterized in yeast, no systematic approach has been performed to study Gα protein coupling of all human serotonin receptors, and the use of current best-performing serotonin GPCRs for biotechnological purposes suffers from low dynamic output ranges (Ehrenworth et al., 2017).

In this paper, we describe the systematic characterization of human serotonin GPCR-mediated biosensing modalities in yeast. Specifically, we characterize signalling in 144 different engineered yeast strains expressing all 12 human serotonin receptors in combination with 12 different Gα protein designs, and furthermore present the characterization of 19 serotonin GPCR polymorphisms mined from the 1,000 Genome Project (1000 Genomes Project Consortium et al., 2015; Spooner et al., 2018). From this, we report serotonin and prucalopride dose-response curves for a total of >30 biosensing designs and apply an optimized biosensing design for high-resolution screening of a yeast strain library engineered to produce serotonin. Collectively, these results provide a resource on chimeric Gα coupling of the human serotonin GPCRs.

## Results

### *Yeast G*α *library screen reveals signalling from activated human serotonin receptors*

In order to systematically investigate the potential to couple any of the 12 human serotonin receptors to the yeast mating pathway, we first mined the Human Protein Atlas database (Uhlén et al., 2015) for tissue- and organ-specific expression patterns of genes encoding the receptors in search of physiological parameters which could be leveraged to confer signalling from these human receptors in a yeast cell.

First, as several of the genes of interest produced splice variants, we selected the corresponding Ensembl transcript IDs **(Supplementary Table S1)** by amino acid similarity to the UniProt canonical sequence. From this analysis it is evident that for each of the serotonin GPCRs maximum expression occurs in different tissues (**Figure 1A**). The 5-HT6, 5-HT2C, 5-HT2A and 5-HT5A receptors express most abundantly in the cerebral cortex, just as 5-HT7 expression is maximal in the parathyroid gland. In female reproductive tissues, 5-HT1B and 5-HT1F express in high levels in the placenta, 5-HT1A and 5-HT1E in the ovaries, and 5-HT2B express in the endometrium and cervix most abundantly. In the gastrointestinal tract, 5-HT1D is most abundantly expressed in the small intestine and duodenum. Similarly, 5-HT4 shows high expression in the small intestine, but comparably lower levels in the rectum, colon, and duodenum. Previously, 5-HT4 has been identified to be highly expressed in the gastrointestinal tract and is a target for drugs for gastrointestinal disorders (Manabe et al., 2010; Wong et al., 2010). Taken together, the 5-HT class of receptors are expressed at different abundances and tissues.

**Figure 1:**
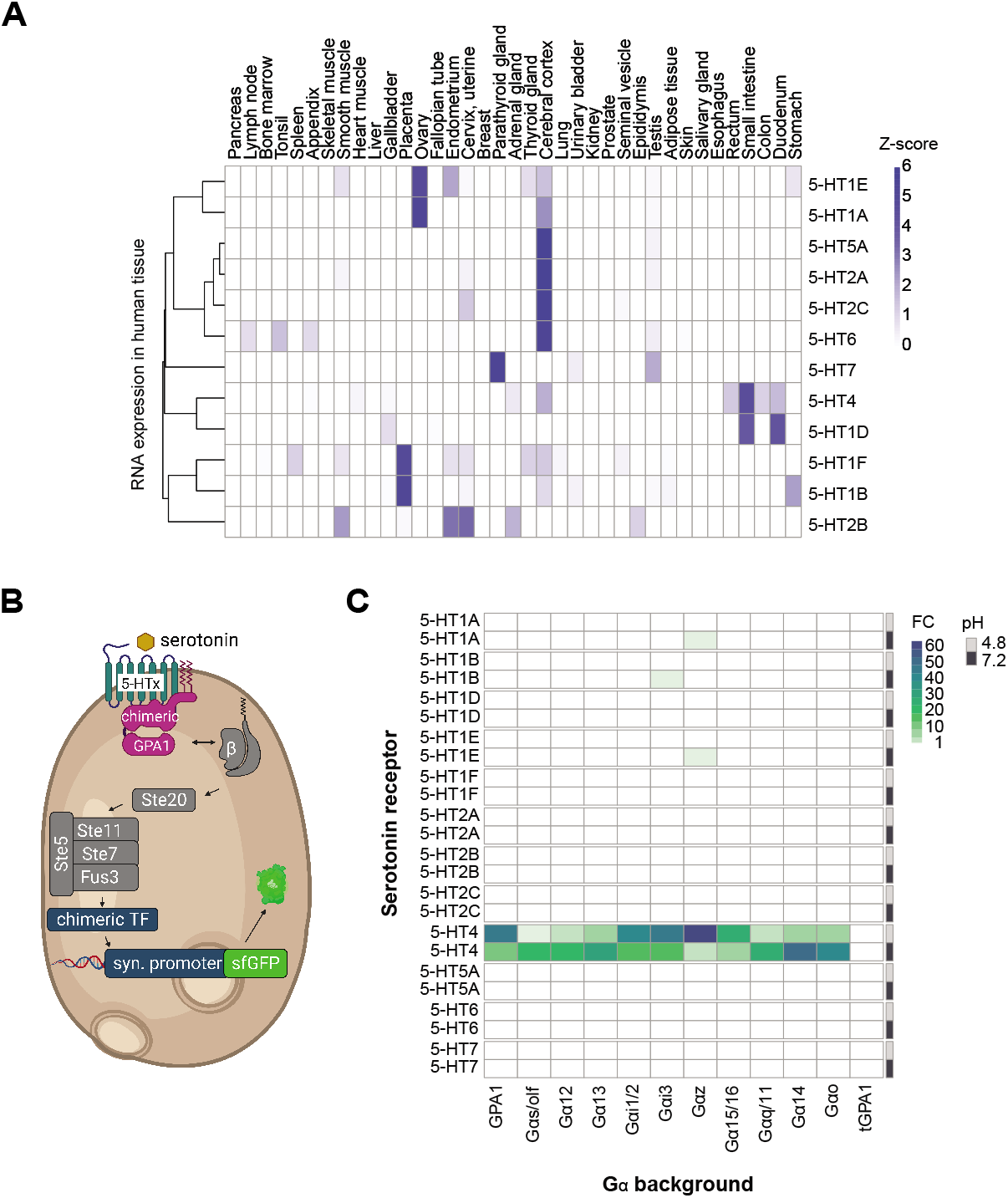
Exploring serotonin GPCR functionality in yeast. **A**) Heatmap of transcripts of serotonin GPCR expression in human tissues and organs from Human Protein Atlas (Uhlén et al., 2015). Color key indicates relative expression levels normalized by row. **B**) Schematic of engineered yeast mating pathway (Shaw et al., 2019), coupled to human serotonin GPCRs. Serotonin binds to 5-HT class of GPCRs, the associated engineered GPA1-based chimeric Gα protein dissociates into one Gα subunit, and a Gβγ dimer to induce the MAP-kinase cascade (Ste5/Ste7/Ste11), which in turn activates a chimeric transcription factor (chimeric TF), binding to an synthetic promoter to enable expression of superfolder green fluorescent protein (sfGFP). **C**) Heatmap of 12 serotonin GPCRs expressed from centromeric plasmids in 12 different Gα background yeast strains. Fold-change shown in color (FC) represents the ratio of fluorescence between induced (100 μM serotonin) and uninduced state (0 μM serotonin). FC values represent the average of triplicate median values sampled by flow cytometry with 10,000 events analyzed for each triplicate in both induced and non-induced conditions. Note that breaks in the color range for 1C are not equidistant for the lower end of the scale to allow for representation of the lower-induced variants.

Next, we performed a combinatorial library screen founded on 12 different Gα protein background strains expressing either a yeast-native GPA1 Gα protein, one of 10 GPA1/Gα chimeras, or a truncated GPA1 (tGPA1) serving as a negative control (Brown et al., 2000; Shaw et al., 2019). Based on this platform, we transformed plasmids containing one of each of the 12 human serotonin GPCRs into the 12 different Gα background strains creating a library of 144 serotonin GPCR:Gα strains. In this setup, successful coupling of a human serotonin GPCRs with the yeast mating pathway will result in the activation of a synthetic transcription factor, which binds to a synthetic promoter to induce expression of sfGFP in the presence of serotonin (**Figure 1B**). As several serotonin GPCRs are highly expressed in both gastrointestinal tissues with lowered pH as well as in the brain (**Figure 1A**), we hypothesized that pH could play a role in receptor expression and/or coupling, and thus we screened the GPCR:Gα library at both pH 4.8 and 7.2 spanning a physiological relevant pH range for both human serum and yeast cultivation medium (**Figure 1C**). At both pHs we cultivated the library in the absence of serotonin and in the presence of 100 μM serotonin and scored relative GFP reporter read-outs following 4 hrs of induction.

From this screen, we found strains expressing 5-HT4 to be activated by serotonin, at both pH 4.8 and 7.2, and in all Gα backgrounds excluding the truncated Gα control (tGPA1) (**Figure 1C**). Fold-inductions for the 5-HT4 receptor at pH 4.8 ranged between 1.8-fold for Gαs/olf coupling and 64-fold for Gαz coupling, followed by 48-fold Gαi3 and 46-fold for yeast-native GPA1. Interestingly, looking at the three highest-induced Gα backgrounds for 5-HT4 at pH 4.8; Gαz, Gαi3 and GPA1, they showed reduced fold-inductions of 4, 17 and 13, respectively when using media at pH 7.2 (**Figure 1C**). In contrast, low-induced 5-HT4:Gα background strains tested at pH 4.8, showed high inductions at pH 7.2 with Gα14 at 51-fold, Gα13 at 33-fold, and Gαq/11 at 26-fold (**Supplementary Table S2, Figure 1C**).

Generally, the absolute induced signal is higher for all strains at pH 7.2 compared to pH 4.8 (**Supplementary Table S2**). The increase in background fluorescence at pH 7.2, rather than a drop in the maximal induced reporter output, is the main reason explaining the overall diminished fold-change for the 3 best performing receptors at pH 4.8. Interestingly, at pH 7.2 many previously poorly activated receptors showed an approximately 10-fold increase in absolute fluorescence levels in the induced state, while the background was only modestly elevated. This is exemplified for the 5-HT4 in the Gα14 background, where total induced reporter gene expression from 25 to 231 is observed while the background fluorescence only increased from 3.03 to 4.56, ultimately shifting the fold-change of 5-HT4 in the Gα14 background from 8.4 to 50.7 when comparing pH 4.8 and 7.2 (**Supplementary Table S2, Figure 1C**). Similar shifts can be observed for previously poorly induced 5-HT4:Gα backgrounds (Gαs/olf,Gα12, Gα13, Gαq/11, Gα14, Gαo) at 4.8, which show foldchanges increased by up to 10-fold at pH 7.2 (**Supplementary Figure S1**, **Supplementary Table S2**, **Figure 1C**).

Furthermore, at pH 7.2, 5-HT1B in the Gαi3 background as well as 5-HT1A in Gαz background showed modest fold-changes of 1.5-fold, while 5-HT1E in the Gαz background reached 1.7 fold-change (**Figure 1C**, **Supplementary Table S2**). In comparison with previous serotonin receptor studies in yeast, 5-HT1A has been shown to couple to Gαo, Gαi2, Gαi3, Gα14, and GPA1, while 5-HT1D has been coupled to Gαi2 and Gαi3 when expressed from high-copy plasmids and using a β-galactosidase or ZnGreen reporter assays (Brown et al., 2000; Nakamura et al., 2015). The observation that 5-HT1A couples to GPA1 was not supported by Ehrenworth *et al*. (Ehrenworth et al., 2017), and neither were we able to detect any changes in reporter output upon serotonin supplementation to strains expressing 5-HT1A, except in the Gαz background (**Figure 1C**). Furthermore, while we could not demonstrate activation of the 5-HT1D receptor at pH 4.8 or 7.2 when the receptors were expressed from single-copy plasmids, our library screen corroborated the recent study by Kapolka *et al*. (Kapolka et al., 2020), showing promiscuity of 5-HT4 in coupling to all chimeric Gα proteins tested.

Taken together, our results show the functionality of 5-HT1A, 5-HT1B, 5-HT1E and 5-HT4 in different chimeric GPA1/Gα backgrounds, with up to 64-fold induction in signalling output. Also, our study illustrates increased background fluorescence at pH 7.2, especially for highly ligand induced Gα protein variants at pH 4.8, and highly increased absolute overall fluorescence for variants poorly induced at pH 7.2.

### Chimeric Gα background impacts EC_50_ and sensitivity of 5-HT4 biosensing

In humans, serotonin receptors are believed to couple and activate inner-cellular responses primarily through the Gas protein (Berumen et al., 2012). To investigate the potential impact different chimeric Gα could have on serotonin sensing in yeast, we next studied the dose-response curves of the 5-HT4 receptor expressed together with the 12 different Gα background strains (**Figure 1B-C**). As genomic integration of serotonin GPCRs shows a more homogenous sensor signal compared to plasmid-based expression of serotonin GPCRs (**Supplementary Figure S2**), 5-HT4 was integrated into the 12 different Gα background strains, and reporter gene outputs obtained with serotonin stimulation between 0.01 - 1,000 μM of serotonin (**Figure 2**). Next, serotonin concentrations yielding the half-maximal reporter output (EC_50_) and sensitivity of serotonin biosensing (Hill coefficient) were obtained. Here, 5-HT4 expressed together with yeast native Gα protein GPA1 showed EC_50_ and Hill coefficient of 49.6 μM and 1.53, respectively (**Figure 2**), while 5-HT4 expressed together with Gαz, Gαi1/2 and Gαi3 all yielded the lowest EC_50_ values of 8.33 μM, 17.17 μM and 35.33 μM, respectively, and Gα12 and Gα14 showed the highest EC_50_ values >200 μM (**Figure 2**). With respect to cooperativity, all designs had Hill coefficients >1, with the highest seen for 5-HT4 expressed with Gαq/11 (2.23), Gα13 (2.07) and Gαo (1.80).

**Figure 2.**
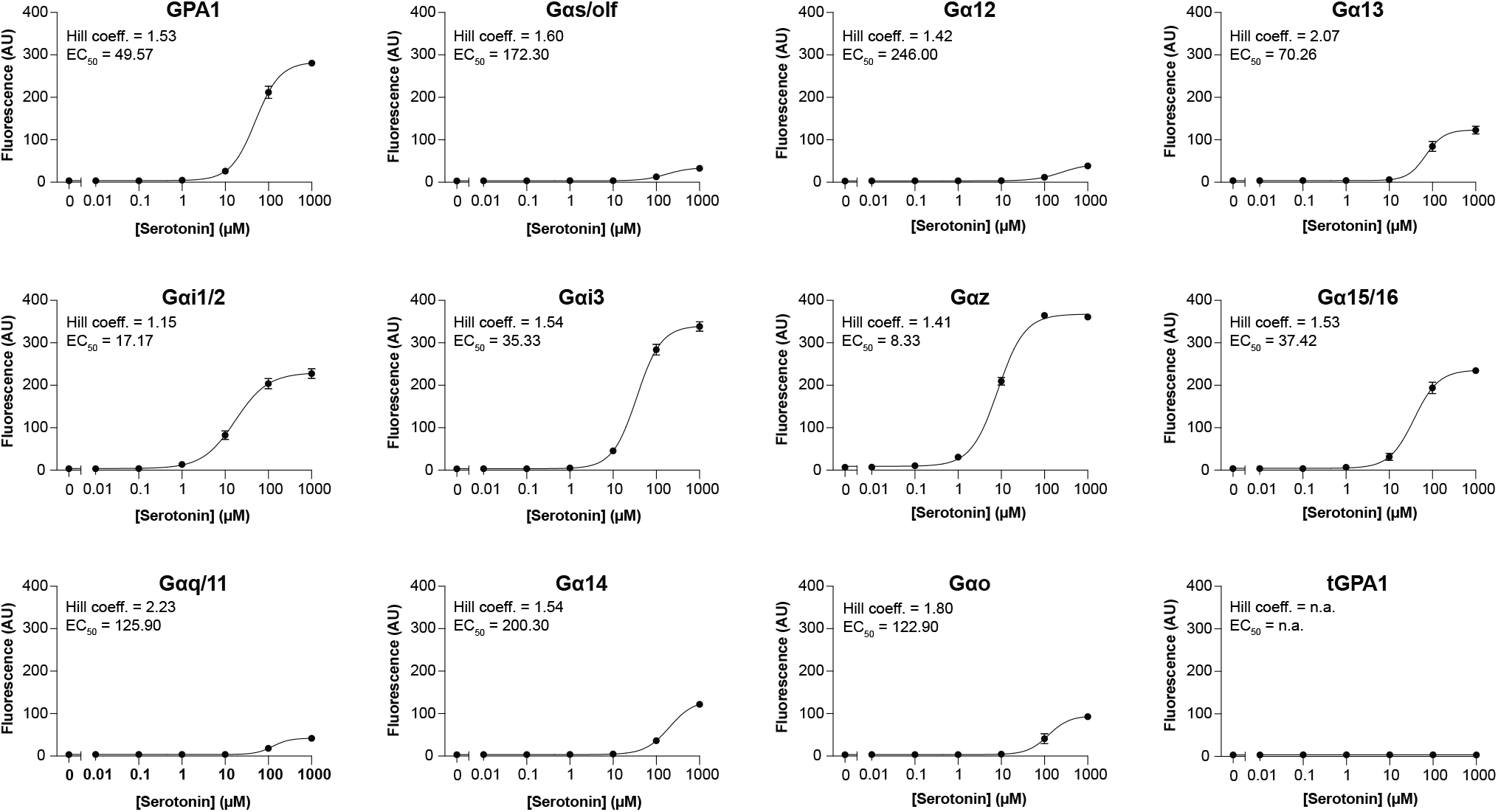
Dose-response curves of 5-HT4 coupling to chimeric Gα proteins. Yeast strains expressing 5-HT4 in combination with 12 different Gα backgrounds, namely yeast GPA1, truncated GPA1 (tGPA1), or any of the 10 different GPA1/Gα chimera (Brown et al., 2000; Shaw et al., 2019). Strains were cultivated in control medium without serotonin, or 0.01 - 1000 μM serotonin, and sfGFP reporter outputs recorded following 4 hrs. All data points represent the median fluorescence intensity of three technical replicates (10,000 events each), of which the mean +/- standard deviation was calculated. Data was fitted to a variable slope four-parameter curve fitting model, from which EC_50_ and Hill slope values were calculated, except for the tGPA1 background strain serving as a negative control. AU = arbitrary units. n.a. = not applicable.

Taken together, serotonin biosensing spanned approximately 25- and 2-fold difference in EC_50_ and sensitivity, respectively, by only using sensor strains with different 5 C-terminal residues of the yeast Gα protein GPA1 changed to C-terminal residues of human Gα proteins.

### A whole-cell biosensing workflow for serotonin

Based on the operational range spanning almost three orders of magnitude, the low EC_50_, and high dynamic range (**Figure 1C & Figure 2**), the chimeric GPA1/Gαz expressed together with 5-HT4 was next chosen as a platform design to explore the possibility of whole-cell biosensing of serotonin produced from yeast. Previously, it was shown that metabolically produced serotonin and melatonin can be sensed using their respective GPCRs (Ehrenworth et al., 2017; Shaw et al., 2019), and also that 5-HT4 could be used as a biosensor to discriminate between reporter outputs from a wild-type yeast and a serotonin-producing yeast, albeit with a modest sensor response of ~2-fold (Ehrenworth et al., 2017). Here, we set out to i) identify key parameters in developing a high-resolution and simple serotonin biosensing workflow using the biosensor based on chimeric GPA1/Gαz expressed together with 5-HT4, and ii) construct a library of variant serotonin-producing yeast strains in order to validate biosensor performance.

Serotonin is produced from L-tryptophan via a 5-hydroxy-tryptophan (5-HTP) intermediate (**Figure 3A**)(Germann et al., 2016). To investigate possible activation of the 5-HT4 sensor by precursor products, the biosensing strain was subjected to L-tryptophan and 5-HTP, as well as serotonin as a positive control over a range of 0.01 μM - 1,000 μM. Activation of the receptor, as inferred by fluorescence output, was only observed for serotonin, confirming the specificity of the 5-HT4 for serotonin over its precursors (**Figure 3B**).

**Figure 3.**
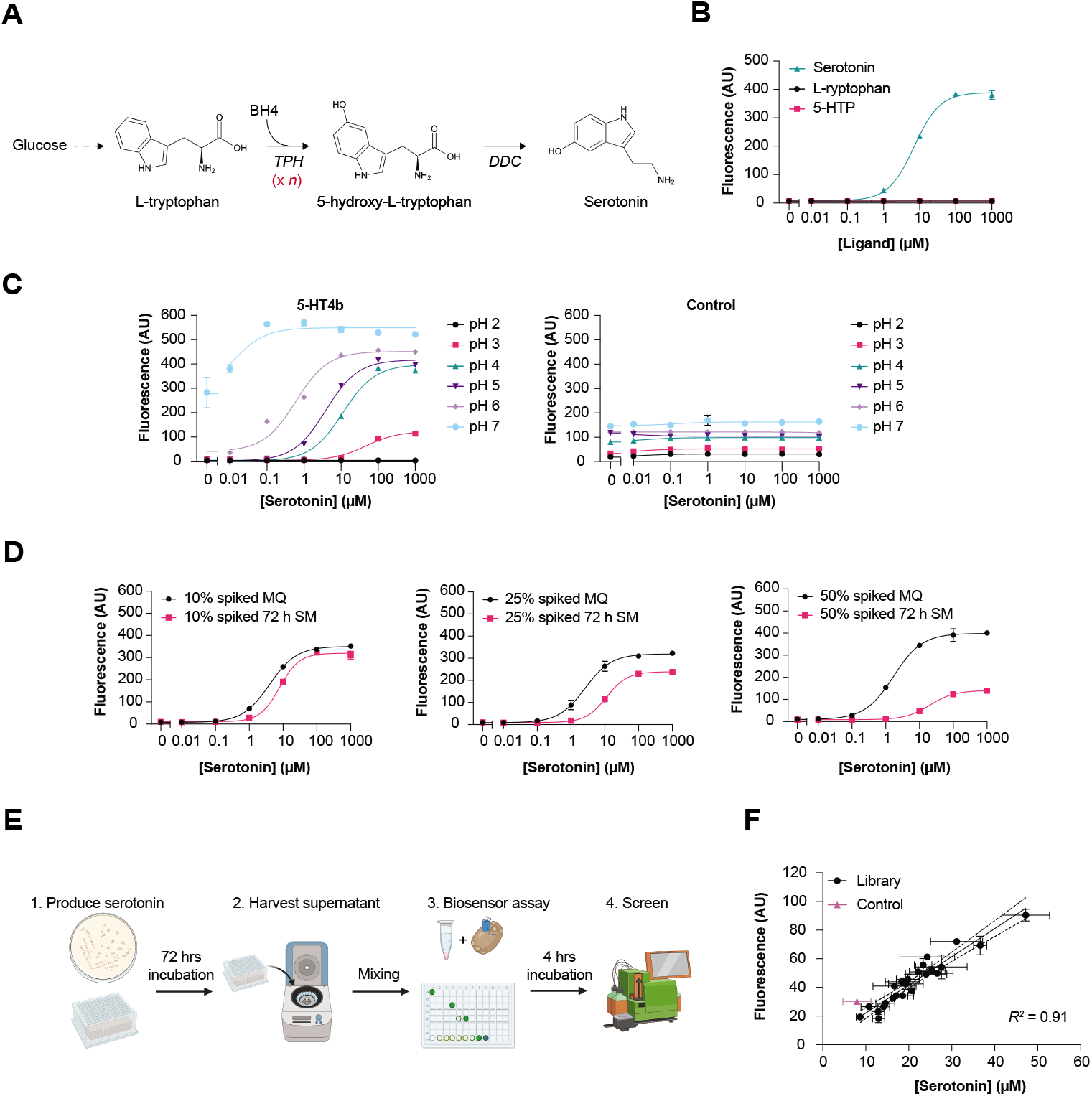
A workflow for semi-throughput characterization of serotonin accumulation in engineered yeast cells. **A)** Serotonin is produced from L-tryptophan and BH4 cofactor via 5-hydroxy-L-tryptophan using tryptophan hydroxylase (TPH) and 5-hydroxy-L-tryptophan decarboxylase (DDC) enzymes. **B**) Dose-response curves of 5-HT4:Gαz sensor strain upon induction with serotonin, L-tryptophan or 5-HTP. **C)** Effect of media with different pH on the Gαz+5-HT4 sensor strain and a constitutively sfGFP expressing yeast base strain, incubated with serotonin **D)** Dose-response curves of adding spiked spent media (72 h SM) or serotonin spiked water (MQ) at different volumetric %-ages added to the sensor strain. **E)** Workflow for sensing yeast-produced serotonin. Serotonin producing cells are incubated for 72 hrs, the supernatant is spun down, added to, and incubated with the sensor strain expressing 5-HT4:Gαz sensor, and sfGFP expression ultimately screened using flow cytometer as a proxy for serotonin production. **F**) Correlation of HPLC-based quantification of serotonin and sfGFP expression via Gαz+5-HT4 of serotonin producing yeast strains carrying multiple copies of TPH (x *n*). Data were fitted with a simple linear regression model. For B) and D), each data point consists of technical triplicates of 10,000 events, for C) 6,500 events were recorded and F) 5,000 events were analysed for each triplicate. For all data panels, the median fluorescence of each triplicate was calculated, and mean ± standard deviation shown.

Next, as we previously observed a strong pH-dependent effect on overall fluorescence and background fluorescence output from yeast strains expressing 5-HT4 together with Gαz (5-HT4:Gαz)(**Figure 1C**, **Supplementary Figure S1, Supplementary Table S2**), we sought to investigate the effect of pH on serotonin dose-response curves over a wider pH range. Consequently, the sensor strain was subjected to media with pH ranging from pH 2 to pH 7. A yeast strain carrying only sfGFP under a *TDH3* promoter served as a control and was subjected to the same pH conditions. Overall, the lowest EC_50_ value is observed at pH 7 (0.01 μM) and EC_50_ values show an inverse proportional relationship with pH (**Figure 3C**). The broadest operational ranges as inferred by changes in sfGFP output are seen for the biosensing strain cultivated at pH 5 and pH 6, spanning from 0.1 to 100 μM, and from 0.01 - 10 μM, respectively. At pH 7, the sensor strain reported changes in serotonin concentrations from 0.01 to 0.1 μM, while at pH 2 no changes in reporter output were observed over the applied range of serotonin concentrations (**Figure 3C**). Of importance, background fluorescence in the absence of serotonin is generally low for all pH conditions tested, with EC_50_ values decreasing by lowering the pH (**Supplementary Table S3**). However, at pH 6 and especially pH 7, background fluorescence increases, as also observed with strains having plasmid-based expression of GPCRs (**Figure 3C**, **Supplementary Figure S1**). Finally, the sfGFP control strain only shows serotonin-independent increase in fluorescence with increasing pH (**Figure 3C**).

In addition to assessing pH effects, and acknowledging that serotonin produced from yeast cells is secreted to the cultivation medium, we tested if adding serotonin in yeast spent media would influence the signalling behaviour of the 5-HT4:Gαz sensor strain. For this purpose, spent media from a yeast base strain (BY4741) cultivated for 72 hrs (72 h SM) was spiked with different concentrations of serotonin. Adding serotonin-spiked SM at 10%, 25% or 50% of the volume in the plate dilution step allowed us to evaluate spent media effects versus a control with serotonin spiked into water (MQ)(**Figure 3D**). At 10%, only a slight decrease in fluorescence was observed between the cultivations with MQ control and 72h SM, as well as a modest increase in the EC_50_ from 4.00 μM to 7.88 μM. At 25% SM, an EC_50_ increase from 2.59 μM to 11.21 μM was observed, while most notably, the EC_50_ increased from 1.70 μM to 20.41 μM when adding the ligand at 50% SM (**Figure 3D, Supplementary Table S4**). Thus, taking into consideration the expected concentration of ligand produced, the ratio at which SM supernatant is added to the medium with the sensor strain should enable a simple biosensing workflow with adjustable EC_50_ values to the application of interest.

Lastly, we applied the workflow to screen a panel of yeast cells engineered to produce different levels of serotonin by randomly integrating variable numbers of expression cassettes for TPH enzyme into genomic Ty2 retrotransposon sites (**Figure 3E-F**)(Germann et al., 2016). Briefly, following random integration of open reading frames for TPH expression in transposable elements of the yeast genome (Germann et al., 2016; Maury et al., 2016), 24 randomly sampled colonies were grown for 72 hrs before harvesting and adding supernatants to the sensor strain, followed by incubation for 4 hrs and measurement of sfGFP fluorescence (**Figure 3E**). In parallel, the supernatants were analysed using HPLC to validate sfGFP reporter output as a proxy for absolute serotonin concentrations in the spent medium.

Taking into consideration acidification of yeast media over prolonged cultivations of strain BY4741 (Murakami et al., 2011), and the observed negative impact of spent media on maximum reporter output (**Figure 3D**), the reporter output from this screen was expected to be diminished compared to the serotonin-spike in titrations (**Figure 2)**. Therefore, for the biosensing workflow, we decided to use synthetic complete medium at pH 4.9, with supernatants from each of the 24 randomly sampled colonies of yeast strains engineered for serotonin production added at 10% volume. While adding spent medium at 10% infers a 10-fold dilution, the biosensor was able to resolve fluorescence outputs in these lower serotonin ranges (**Figure 3F**). From plotting biosensor fluorescence outputs against serotonin quantification as inferred from HPLC, a linear model fitted HPLC-measured serotonin concentrations and biosensor fluorescence from the 24 sampled strains (*R*^2^ = 0.91)(**Figure 3F**).

In summary, the engineered 5-HT4:Gαz biosensing strain specifically senses serotonin, and can reliably detect serotonin in a facile and easy-adjustable (e.g. pH and spent medium) workflow compatible with high-throughput screening of libraries of yeast cells engineered to produce serotonin.

### Characterization of human 5-HT4 variants in yeast

GPCR single-nucleotide polymorphisms (SNPs) are known to impact EC_50_ and agonist sensitivities in humans (Marti-Solano et al., 2020), and human variants of 5HT1a and MOR1 expressed in yeast have previously shown to reproduce Gα-dependent sensitivities to serotonin and morphine, respectively, as reported from mammalian cells (Bean et al., 2021; Nakamura et al., 2015).

Based on the biosensing platform developed in this study we next sought to examine the canonical isoform b of human 5-HT4, in comparison to human receptor variants or ‘protein haplotypes’ sourced from the 1,000 Genomes Project using the Haplosaurus tool browser via Ensembl (1000 Genomes Project Consortium et al., 2015; Spooner et al., 2018). From this data mining, 20 5-HT4 receptor variants were identified, of which 19 were cloned into yeast (**Figure 4A**). Nine of the tested variants were in the intracellular loops (including two on residue 137), four were located in the transmembrane domains, four on the C-terminus, one in the extracellular loop, and one in the N-terminus **(Figure 4A)**.

**Figure 4.**
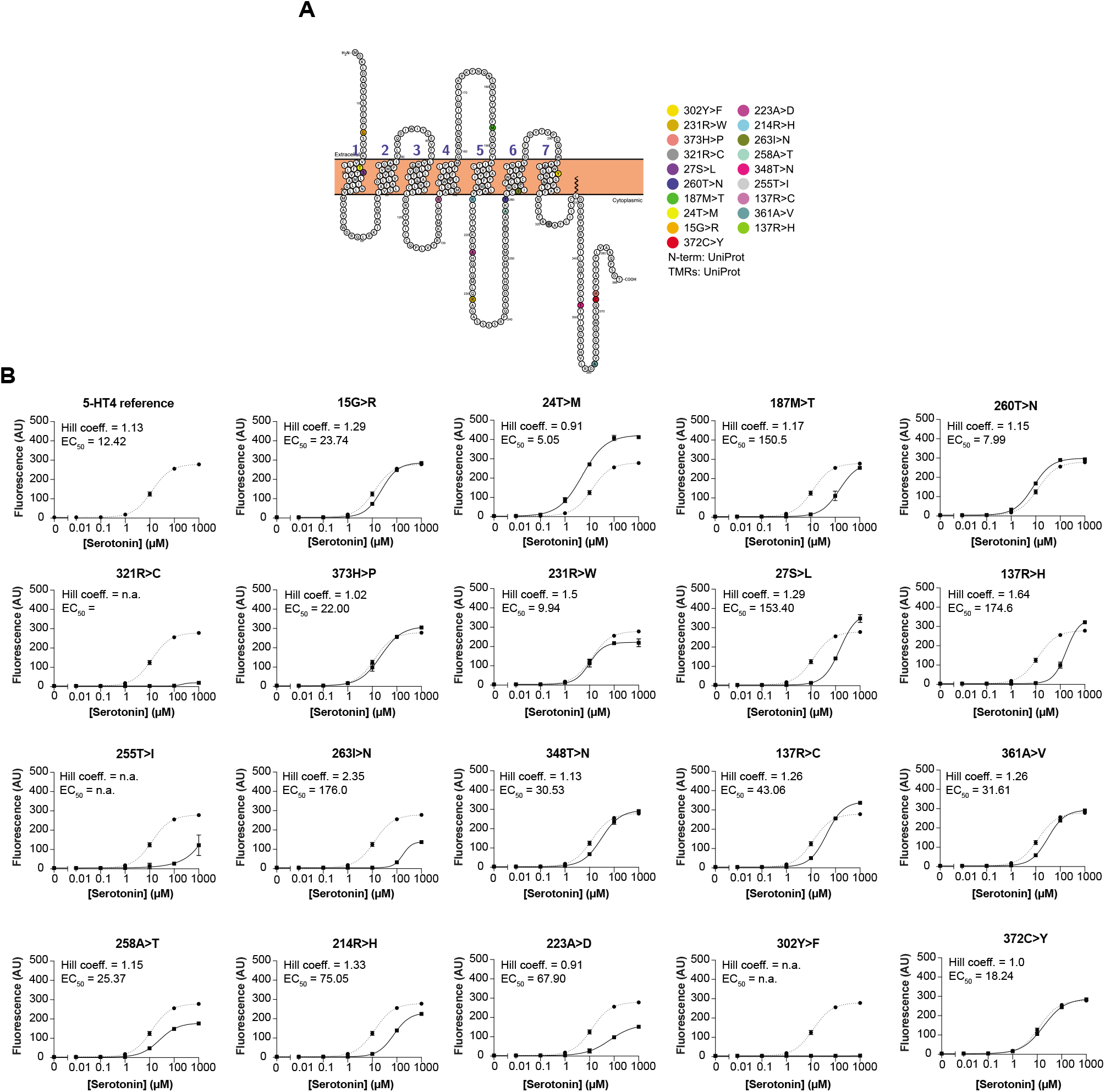
Characterization of 5-HT4 serotonin GPCR variants from human genomes. **A**) Snake-plot of 5-HT4 isoform b mutational landscape showing location of analyzed variants on the protein. All screened variants had a single mutated amino acid. Two different variants were screened separately on residue 137. **B**) Dose responses for the 5-HT4b isoform reference receptor and 19 variants in GPA1 background. Yeast strains expressing single-amino acid variant GPCRs were induced with 0.01 μM - 1,000 μM serotonin in addition to a non-induced “0” control. sfGFP was measured following 4 hrs of incubation with serotonin. All data points represent median fluorescence intensity of three technical replicates (10,000 events each), from which the mean +/- the standard deviation was calculated. AU = arbitrary units, n.a. = not applicable. Data was fitted to a variable slope four-parameter curve fitting model, from which EC_50_ and Hill slope values were calculated. Dose-response curves show the tested variant (solid line) and the reference receptor (dashed line).

Filtered data from Ensembl’s Haplosaurus tool shows the distribution of variants across five different populations: African, American, East Asian, European and South Asian (**Table 1**). From this distribution, there are 218 non-reference genomes present in the dataset, showing different frequencies of variants in different populations (**Table 1**). Of these variants, some were computationally predicted to have deleterious or possibly damaging effects on receptor function (“D”, **Table 1**). Because of this, we decided to introduce the receptor variants into the GPA1 background, as this background has a similar dynamic output range, but higher EC_50_ compared to the Gαz background (**Figure 2**). Next, for the 5-HT4 dose-response study (**Figure 2**), receptor variants were assayed for serotonin responsiveness from 0.01 - 1,000 μM serotonin, in addition to non-induced control without supplemented serotonin. EC_50_ values and Hill coefficients ranged from 5.05 μM to 203.7 μM, and 0.67 to 3.20, respectively (**Figure 4B**). The 5-HT4 isoform b reference strain had an EC_50_ of 12.42 μM and Hill coefficient of 1.13. while variants 24T>M (5.05 μM), 260T>N (7.99 μM), and 231R>W (9.94 μM) resulted in decreased EC_50_ values. Additionally, the 24T>M variant had an increased operational range as compared to reference **(Figure 4B)**. Oppositely, the variants 372C>Y, 15G>R, 373H>P, 348T>N, and 361A>V variants resulted in minor increases in EC_50_ values (1824 - 31.61 μM), while variants 27S>L, 321R>C, 137R>H, and 263I>N variants resulted in notably higher EC_50_ values (153.4 - 176.0 μM) as compared to the reference receptor. Interestingly, the 137R>H variant had a considerably higher EC_50_ value than the 137R>C variant (174.6 vs 43.06 μM) despite their shared residue location. For variants 302Y>F and 255 T>I no EC_50_ or Hill coefficient could be calculated due to their high detection limits and almost complete loss of function, respectively. Of the variants located in the transmembrane domains, 24T>M decreased the EC_50_ and increased the operational range, whereas 27S>L, and 263I>N resulted in EC_50_ values over 100 μM (**Figure 4A-B**).

**Table 1.** Demographic distribution of human 5-HT4 non-synonymous polymorphisms. Demographic distribution of 5-HT4b haplotypes adapted from 1,000 Genomes data using the Haplosaurus tool (Spooner et al., 2018). Haplosaurus annotated variants with a ‘D’ in the ‘Flags’ column that are flagged as being deleterious (SIFT) or probably damaging (PolyPhen-2). Frequency data comes from the GRCh38 reference genome from the Genome reference Consortium using data from the 1,000 Genomes Project phase 3.

| Protein Haplotype/Variant | Flags | Count | African | American | East Asian | European | South Asian |
| --- | --- | --- | --- | --- | --- | --- | --- |
| Reference |  | 5799 | 1208 | 688 | 1004 | 1003 | 970 |
| 372C>Y |  | 158 | 95 | 4 | 0 | 0 | 0 |
| 15G>R |  | 26 | 13 | 0 | 0 | 0 | 0 |
| 24T>M |  | 9 | 2 | 1 | 0 | 0 | 0 |
| 187M>T | D | 4 | 1 | 0 | 0 | 0 | 0 |
| 260T>N | D | 3 | 0 | 0 | 0 | 0 | 3 |
| 27S>L |  | 2 | 0 | 0 | 0 | 1 | 1 |
| 321R>C | D | 2 | 0 | 0 | 2 | 0 | 0 |
| 373H>P |  | 2 | 0 | 0 | 0 | 0 | 0 |
| 231R>W | D | 2 | 0 | 1 | 0 | 0 | 0 |
| 302Y>F | D | 1 | 0 | 0 | 0 | 0 | 1 |
| 137R>H | D | 1 | 0 | 0 | 0 | 1 | 0 |
| 361A>V |  | 1 | 1 | 0 | 0 | 0 | 0 |
| 137R>C | D | 1 | 0 | 0 | 0 | 1 | 0 |
| 255T>I | D | 1 | 1 | 0 | 0 | 0 | 0 |
| 348T>N |  | 1 | 0 | 0 | 0 | 0 | 1 |
| 258A>T | D | 1 | 1 | 0 | 0 | 0 | 0 |
| 263I>N | D | 1 | 0 | 0 | 1 | 0 | 0 |
| 214R>H | D | 1 | 0 | 0 | 0 | 0 | 1 |
| 223A>D | D | 1 | 0 | 0 | 0 | 0 | 1 |

To further investigate the impact of the 5-HT4 variants, we tested our variant library with the 5-HT4 agonist, prucalopride succinate. Prucalopride is a high affinity, highly selective 5-HT4 receptor agonist designed to treat gastrointestinal disorders such as constipation (Briejer et al., 2001). Dose responses with prucalopride revealed a lower operational range with a reference EC_50_ of 87.48 μM and a Hill coefficient of 0.88, while the 24T>M variant maintained the increased dynamic range as observed with serotonin and a low EC_50_ of 8.92 μM and Hill coefficient of 1.17 (**Supplementary Figure S3**). Additionally, of note, 302Y>F maintained the loss-of-function, just as receptor variants 137R>H, 137R>C, 223A>D, 255T>I, 263I>N, 321R>C also showed lower or loss-of-function in the presence of prucalopride.

Ultimately, this study enabled reporting of dose-response parameters for 19 5-HT4 variants found in human populations spanning five demographic regions with serotonin and prucalopride. Importantly, in yeast, variants stimulated with serotonin displayed large differences in EC_50_ values (5.05 - 203.7 μM), cooperativity as inferred from Hill coefficients, as well as maximum reporter output, compared to the reference receptor.

## Discussion

Here we show a combinatorial serotonin GPCR:Gα library screened at different pHs, and find 5-HT4, 5-HT1A, 5-HT1B, and 5-HT1E functional in yeast. In-depth characterization of serotonin-responsiveness of 5-HT4 receptors expressed in the different Gα backgrounds show EC_50_ values ranging from 8 - 250 μM, almost 2-fold changes in cooperativity, and up to 64-fold dynamic output ranges. We also present a high-resolution workflow sensing microbially produced serotonin and show that pH in spent cultivation medium strongly influences GPCR-mediated reporter outputs, allowing for simple adjustments of the sensing workflow according to concentration range of serotonin. Importantly, measuring microbially produced serotonin, the biosensor output strongly correlates with HPLC-measured serotonin concentrations, enabling finding best-producing strains at ease. Lastly, in addition to Gα, pH and spent medium ratio, from a library of 19 5-HT4 receptor variants we also demonstrate that single residue polymorphisms in serotonin GPCRs impact key reporter output parameters like EC_50_, cooperativity, agonist specificity, and maximum output. Taken together these findings enable rational tuning and further characterization of serotonin biosensing modalities.

Important findings from this study are the receptor functionality in yeast for 5-HT4 in all Gα protein backgrounds, and the modest fold-inductions (i.e. 1.5 - 1.7) for 5-HT1B in the Gαi3 background, 5-HT1A in Gαz background, and 5-HT1E in the Gαz background. In agreement with Ehrenworth *et al*. (Ehrenworth et al., 2017) we found that, when coupled to yeast-native GPA1, the 5-HT1A, 5-HT1D 5-HT2B, 5-HT5A and 5-HT6 receptors were nonfunctional at both pHs tested in this study. While our findings largely abrogates recent findings related to Gα coupling and 5-HT receptor subtype, Brown *et al*., showed activation of 5-HT1A in chimeric and wild-type GPA1 Gα proteins (Brown et al., 2000). In our study, we were not able to demonstrate activation of the 5-HT1D receptor, and also not 5-HT1A in the GPA1 and Gαi3 background previously reported (Brown et al., 2000; Nakamura et al., 2015). Likewise, previously, Yasi *et al*. expressed 5-HT4 in *S. cerevisiae*, and when assayed with serotonin and prucalopride they found that serotonin-activated receptors had a higher EC_50_ than prucalopride (Yasi et al., 2019), whereas our study found 5-HT4 to have higher EC_50_ for serotonin compared to prucalopride. Lastly, during preparation of this manuscript, pH-dependent coupling of 5-HT4 for all 10 chimeric Gα was reported, in what was referred to as proton-gated coincidence detection (Kapolka et al., 2021). Interestingly, while Kapolka *et al*. observed 5-HT4 to signal through all Gα proteins at pH 7 and only Gαz and Gαi1/2 at pH 5, we observed coupling to all tested Gα proteins at both pH 4.8 and 7. Some of these divergent findings are likely attributable to different genetic backgrounds of sensor strains, the mode of expression (i.e. low copy plasmid, high copy plasmid, integration), and/or differences in reporters used (i.e. sfGFP, mTq2, ZsGreen, luciferase or β-galactosidase). Ultimately, different assay and strain designs impede direct comparison between studies calling for adoption of assay standards and protocols in regard to cultivation medium, timescales and background strains, when characterizing GPCRs in heterologous hosts like yeast.

While 5-HT1A, 5-HT1B, 5-HT1E or 5-HT4 were demonstrated functional in this study, reasons for non-functionality of the other 8 serotonin GPCRs in yeast could be manifold. Key points to be considered for future mitigations are endoplasmic reticulum processing issues, improper membrane localization, lack of coupling receptors to Gα proteins, and suboptimal sterol environment compared to native human host cells. While adding signaling sequences seems to yield mixed results (Dirnberger and Seuwen, 2007; Fukutani et al., 2012), changing the lipid composition from yeast native ergosterol to sterol synthesis has yielded promising results, shown in the activation of previously non-functional opioid receptors in yeast (Bean et al., 2021). Likewise, varying tissue and organ expression (**Figure 1C**), coincidence detection (Kapolka et al., 2021), and even biased signaling (Choudhary and Loewen, 2016), affect GPCR signaling, and should be considered for further investigation of 5-HT receptor signaling in yeast.

Furthermore, we demonstrated the impact of 19 different polymorphisms of 5-HT4 found in the 1,000 Human Genomes Project on signaling behavior, and compared them to the reference 5-HT4 sequence (1000 Genomes Project Consortium et al., 2015). Previously, a few 5-HT4 variants have been studied in mammalian heterologous systems. For instance, when expressed in 5-HT4 isoform g, 302Y>F decreased the affinity of GR113808 to the receptor by 13-fold but did not affect serotonin induced activity, though it decreased receptor expression (Rivail et al., 2004; Vass et al., 2019). Here, in the *S. cerevisiae* platform, the variant 302Y>F results in a loss of function receptor in the presence of serotonin and prucalopride **(Figure 4, Supplementary Figure S3)**. Furthermore, and of pharmacokinetic relevance, it has been hypothesized that residue 302Y is an important stabilizing residue for agonist and antagonist binding to the 5-HT4 receptor (Mialet et al., 2000; Rivail et al., 2004). From the Haplosaurus tool (Spooner et al., 2018), we observed that the South Asian individual was heterozygous for the 302Y>F variant, which could have implications to responsiveness of 5-HT4 targeting drugs. While we did not study the binding affinity of the 302Y>F variant or its expression in the *S. cerevisiae* platform as performed by Rivail *et al*. (Rivail et al., 2004), our findings highlight that residue 302Y is important for serotonin signaling via 5-HT4. While the yeast platform cannot be 1:1 paralleled to human cells and GPCR signaling, the difference in activity and EC_50_ values in the receptor variants could indicate important shifts in receptor activity and expression.

For future directions, we envision the study of polymorphisms of receptor variants for better sequence- and structure-guided engineering of GPCRs to flourish. As demonstrated herein, changes of one amino-acid in a receptor can vastly influence the signaling behavior of GPCR receptors, and we highlight yeast as a relevant chassis for mutational studies coupled to machine learning approaches, ultimately enabling better understanding of GPCR specificity, expression, pharmacokinetic properties, and finally allow for a more targeted engineering of advanced biosensors Additionally, establishing yeast as a platform to study mammalian receptor polymorphisms could allow for a high-throughput platform for flagging possible variant-related drug activity impacts, and thus serve a purpose in hit-to-lead drug discovery regimes. In terms of human health, such findings should then further extent yeast GPCR biosensing to real-life applications, as recently shown for GPCR-based feedback-regulation loop in an engineered probiotic yeast *Saccharomyces boulardii* for the destruction of extracellular ATP in mice guts (Scott et al., 2021; Shaw et al., 2019). We anticipate that such opportunities will be explored further and form the basis for many more GPCR-based biosensors to be developed and applied in the near future.

## Acknowledgements

This study was supported by Novo Nordisk Foundation Copenhagen Bioscience Ph.D. grant No. NNF19CC0035454 and NNF20SA0035588 and Novo Nordisk Foundation Center for Biosustainability grant number NNF20CC0035580. Authors also would like to thank Luke W. Johnston for help with R scripts, Marcus Deichman for fruitful discussions, as well as Lars Schrübbers for technical assistance related to analytical serotonin measurements.

## Author Contributions

BL, EDJ and MKJ conceived the study. BL and EEH-S performed all the experiments and performed all the data analysis. BL, EEH-S, EDJ, TJ and CNJ designed and constructed all plasmids and strains. BL, EEH-S and MKJ wrote the manuscript. All authors approved the manuscript.

## Declaration of Interests

The authors declare no competing interests.

## Method Details

### Cultivation of E.Coli

Chemically competent *Escherichia coli* DH5α strain was used for plasmid propagation and cloning. E.Coli strains were grown in 2xYT media, supplemented with 100 μg/mL ampicillin at 250 rpm.

### Cultivation of yeast

*Saccharomyces cerevisiae* BY4741 strains described in this study were grown in synthetic complete media (6.7 g/L yeast nitrogen base without amino acids with appropriate drop-out medium supplement) with appropriate auxotrophic selection, supplemented with 20 g/L glucose. Yeast in pre-culture tubes was grown at 30 °C and 250 rpm, while incubation in 96-well deep well plates took place at at 30 °C and 300 rpm. The complete list of all synthetic genes used can be found in **Supplementary Table S5.**

### GPCR and 5-HT4b variant sourcing

The protein sequence of GPCRs was selected on uniprot.com and translated into nucleic acid sequence using the EMBOSS Backtranseq tool (Madeira et al., 2019; UniProt Consortium, 2021) and ordered as biobricks via Twist Bioscience.

For human 5-HT4b variants The identified transcript for human 5HT4b (ENST00000377888) was identified on Ensembl through the International Genome Sample Resource database to find information on global population variants and distribution (1000 Genomes Project Consortium et al., 2015). Single amino acid variants and their population frequency were identified through the Ensembl genome browser using the Haplosaurus tool for the previously specified transcript ID (Spooner et al. 2018). Due to the codon-optimized 5HT4b receptor, variants were designed with the amino acid variation of interest irrespective of base pair changes. Of the 20 5-HT4b single amino acid variants listed on the Haplosaurus protein-haplotype browser on Ensembl, 19 were tested in S. *cerevisiae* due to a cloning issue of one of the variants.

### Plasmid construction and transformation into E. Coli

All plasmids in this study were cloned using USER™ (uracil specific excision reagent) cloning (New England Biolabs) and the EasyClone method (Jessop-Fabre et al., 2016). Genetic parts for assembly into plasmids and USER™ vector plasmids were amplified using PhusionU polymerase (Thermo Fisher Scientific). Plasmids containing the GPCRs had a Kozak sequence (AAAACA) in front of the start codon of the receptor. Synthetic genes were ordered from TWIST Bioscience, custom oligos were ordered from IDT or used from previous publications (Jakočiūnas et al. 2015). The complete list of all gBlocks and plasmids and used can be found in **Supplementary Table S5, S6, and S7.** Plasmids were transformed into chemically competent DH5α strain by heat-shocking for 45 seconds at 42 °C and recovered on LB plates supplemented with 100 μg/mL Ampicillin.

### Yeast transformations

Plasmids and linear DNA parts for integration were transformed into yeast using the lithium acetate/single-stranded carrier DNA/PEG method (Gietz and Schiestl, 2007).

### Construction of yeast strains

For plasmid based GPCR expression, library strains were constructed by transforming plasmids containing the respective serotonin GPCR under a *CCW12* promoter, and a *HIS3* marker. Plasmids were transformed into yeast strains yWS2261-yWS2272, representing optimized sensor strains with different Gα protein backgrounds (Shaw et al., 2019). Transformed yeast cells were selected on SC-HIS plates.

For integration of GPCR into the yeast genome, plasmids with overlap to genomic sites as described by Jessop-Fabre *et al*. were engineered to contain the respective genetic sequence to be integrated (serotonin GPCR, 5-HT4(b) variant) (Jessop-Fabre et al. 2016). Plasmids were NotI digested for 4 hrs, the NotI enzyme was heat-inactivated and linear fragments were integrated into yeast genomic sites with the help of a Cas9 plasmid and a gRNA plasmid targeting the respective integration site (Jessop-Fabre et al., 2016).

The serotonin production strains were constructed as described previously (Germann et al., 2016). Plasmid pCfB9221 containing enzymes HsDDC and SmTPH was previously constructed and integrated into XI-3 (Germann et al., 2016). Cofactor enzymes RnPTS and RnSPR, as well as PaPCBD1 and RnDHPR were cloned on 2 plasmids and integrated into EasyClone sites X-4 and XII-4, respectively. Additionally, plasmid p2772, constructed by (Germann et al., 2016), containing SmTPH with overlap to TY2 sites, was integrated in the yeast TY2 sites.

The complete list of all yeast strains constructed can be found in **Supplementary Table S8.**

### Biosensor assay

Yeast strains were freshly plated and grown on SC plates with respective auxotrophy if required. On day 1, a single colony of a sensing strain was inoculated in SC media with the respective auxotrophy if needed and grown for 24 hrs. On day 2, the culture was diluted 1:100 in SC media with or without auxotrophy and grown for 16 hrs. On day 3, the culture was diluted 1:50 in SC media with or without auxotrophy and grown for two hrs. In 96-well flat-bottom plate, 20 uL of ligand dissolved in either milliQ water, serotonin-producing yeast supernatant or spent media (SigmaAldrich: Serotonin hydrochloride H9523-100MG, L-tryptophan T0254-1G, 5-hydroxy-L-tryptophan H9772-1G) was combined with 180 μL of culture of a sensing strain, except for the experiment in **Figure 3D**, where additionally ratios of 50 uL:150 μL and 100 μL:100 uL ligand:biosensor were used. Note that different serotonin concentrations (that means 10,000 μM, 4,000 μM and 2,000 μM that were added at 10%, 25% or 50% volume, resulting in 1000 μM in-plate-concentration in the highest dilution step. The 96-well plate was covered with a PCR foil and incubated for another 4 hrs at 30 °C and 300 rpm. Plates were chilled at 4 °C until ready for flow cytometry analysis. The same experimental set-up was used for the dose response curves with prucalopride succinate dissolved in milliQ water (SigmaAldrich: Prucalopride succinate AMBH2D6FB305-100MG). pH of the SC-media ranged between 4.7 and 5.3 during all experiments apart from the pH experiment (**Figure 3C**).

### Flow cytometry analysis

Flow cytometry analysis was performed on the Miltenyi MACSQuant VYB, using medium mixing and fast running mode. 10,000 events were recorded for each well analyzed, unless otherwise specified. Cells were gated for singlets in exponential phase, and 10,000 events within the singlet gate were recorded for each well analyzed, apart from Figure 3C, where 6,500 events were recorded, and Figure 3F, where 5000 events were analyzed.

### Serotonin production in yeast

Serotonin producing strains were inoculated from a single colony and grown for 16 hrs in SC-URA media (pH 4.9). Cultures were diluted 1:50 and grown for 72 hrs in SC-URA media in 96-well deep well culture plates. Supernatants were harvested by spinning at 4000 rpm for 5 min. Supernatants were carefully transferred into 96-well plates, covered with aluminum foil and stored at −80 °C for flow cytometry or HPLC analysis.

### HPLC analysis of serotonin production strains

Analysis of serotonin was done on the Thermo Scientific™ UltiMate™ 3000. Solvent A was 0.05% acetic acid, solvent B Acetonitrile. The column used was the Agilent Zorbax C18 4.6×100mm 30l5-Micron with a Phenomenex AFO-8497 filter. The solvent gradient can was:

| Time | Flow<br>[ml/min] | %A | %B |
| --- | --- | --- | --- |
| 0 | 1.000 | 95.0 | 5.0 |
| 1.5 | 1.000 | 88.0 | 12.0 |
| 2.5 | 1.000 | 88.0 | 12.0 |
| 4.5 | 1.000 | 70.0 | 30.0 |
| 5.5 | 1.000 | 70.0 | 30.0 |
| 8.0 | 1.000 | 30.0 | 70.0 |
| 9.0 | 1.000 | 30.0 | 70.0 |
| 9.5 | 1.000 | 95.0 | 5.0 |
| 11.0 | end | end |  |

HPLC values were determined according to two standard curves between 0 and 100 uM serotonin hydrochloride in MQ, one run before and one after the samples between 0 and 100 μM serotonin hydrochloride in MQ using the using Chromeleon™ Chromatography Data System (CDS) Software.

### Data analysis

Data analysis was done in R programming language usingR studio, with customised R scripts, making use of the tidyverse, flowCore and pheatmap packages (Team 2020; RStudio Team 2020; Wickham et al. 2019; Hahne et al. 2009; Kolde 2017).

For dose response curves with flow cytometry data and the heatmap in Figure 1C, data were always obtained in triplicate and median fluorescence values of each triplicate (consisting of min. 5000 events as described in the biosensor assay section) were calculated, of which the mean and standard deviation was calculated. Mean and standard deviation were exported to GraphPadPrism to create dose-response curves. Curve fitting was done in GraphPadPrims with the variable 4 parameter model, apart from Figure 3C, in which curves were fitted with the three-parameter model. Only curve fits >0.9 were considered, if lower, or EC_50_ and hillslopes could not be calculated, “n.a.” is stated. For Figure 3F, simpler linear regression in GraphPadPrism was used (GraphPad Software, San Diego, California USA). For Figure 1C, fold-changes were calculated by dividing the induced state value fluorescence intensity by the uninduced state fluorescence after calculating the mean of 3 median values as described previously. Sensor strains with fold-change values >1.4 were considered functional.

HPLC data were analysed using Chromeleon™ Chromatography Data System (CDS) Software. The snake plot was constructed using Protter (Omasits et al., 2014) and with structure 5HT4R_HUMAN imported from UniProt (UniProt Consortium 2021).

For the transcript heatmap in **Fig. 1A** tissue isoform-RNA data was sourced from the Human Protein Atlas database (Uhlén et al. 2015). Data is available from proteinatlas.org, version 20.1. Several of the genes of interest produced splice variants so corresponding Ensembl transcript IDs were selected by amino acid similarity to the UniProt canonical sequence. In the case where several splice variants matched the canonical sequence, transcript levels were compared in GTEx and the transcript with higher tissue expression was selected (Carithers et al. 2015). Non-tissue samples were filtered out from the dataset. Heatmaps were made with the R package ‘pheatmap’ and normalized by row (Hahne et al., 2009; Kolde, 2017; Team, 2020; Wickham et al., 2019). Transcript IDs for receptors can be found in **Supplementary Table S1**.

All scripts for data analysis can be found at https://github.com/betlen/serosense.

## Notes

https://github.com/betlen/serosense

### Competing Interest Statement

The authors have declared no competing interest.

