## Supplemental Information for "Serotonin GPCR-based biosensing modalities in yeast"

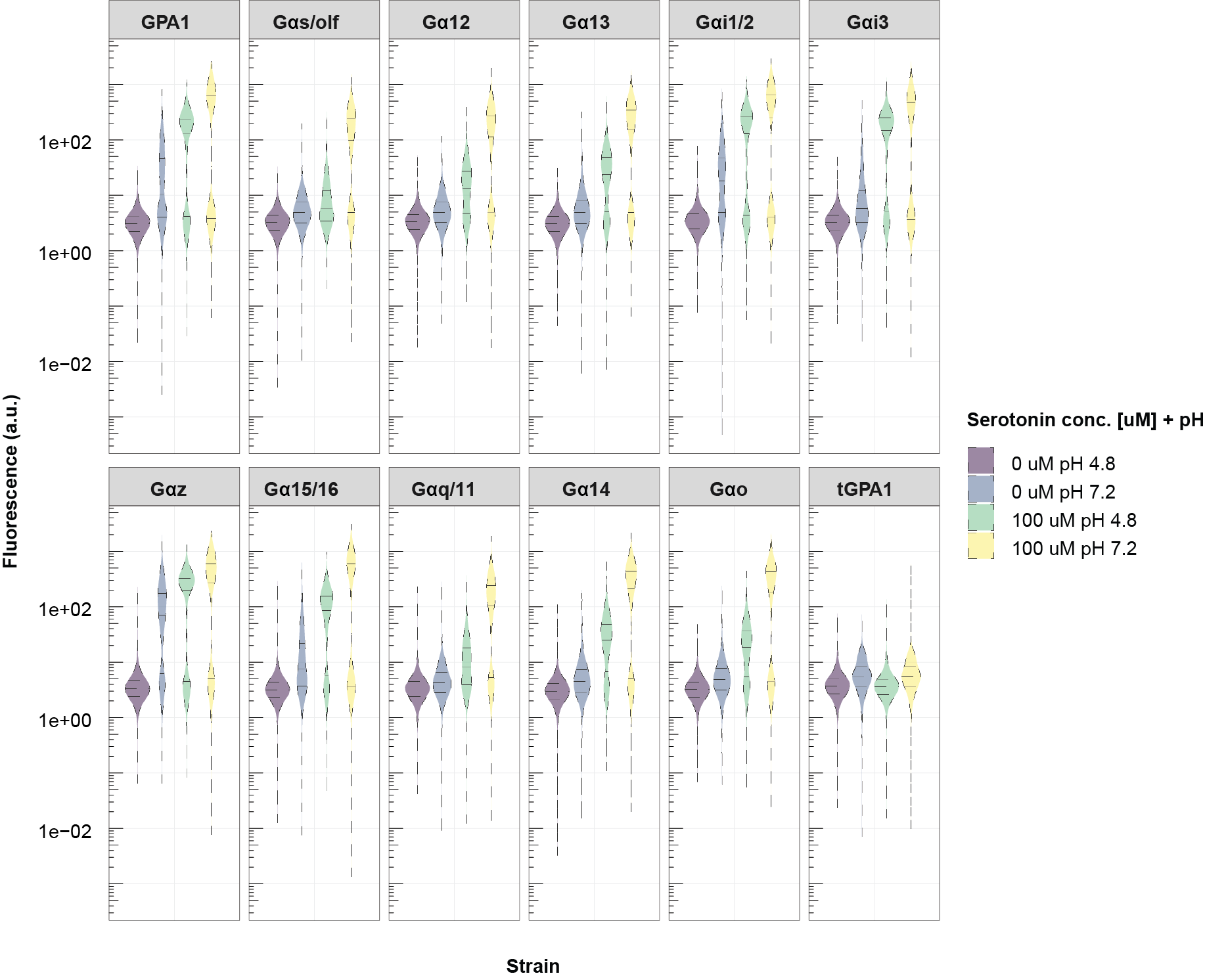

**Supplementary Figure S1.** 12 yeast strains with different Gα protein backgrounds, transformed with 5-HT4 plasmids. Strains were subjected to 100 μM and 0 μM serotonin with the assay run either at 4.8 or 7.2. Each violin plot represents 10,000 events.

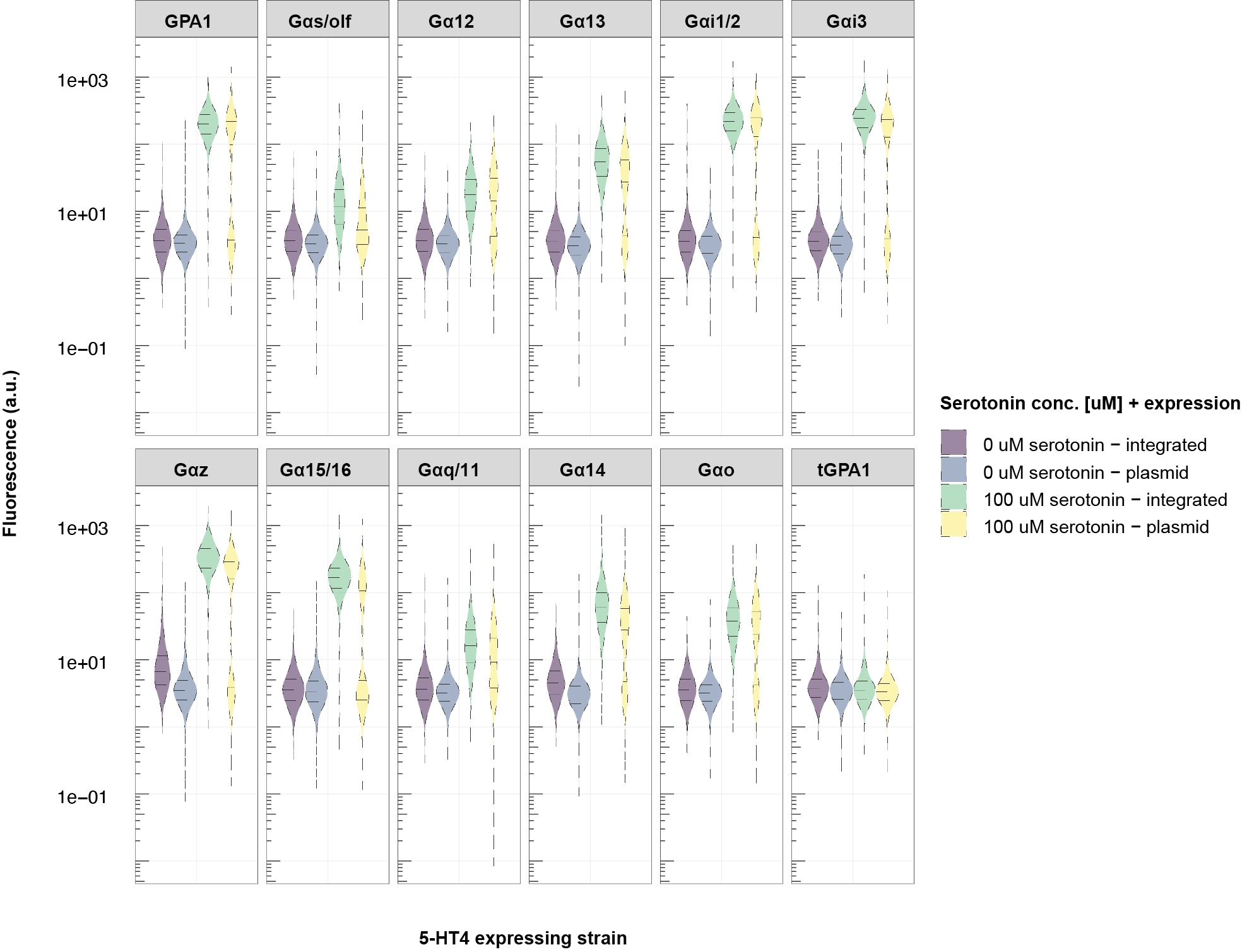

**Figure S2.** Fluorescence of yeast biosensors strains with different Gα protein backgrounds, either having 5-HT4 integrated (purple and green) genomically or expressed from a plasmid (blue or yellow). Strains were subjected to 100 μM and 0 μM serotonin. Each violin plot represents 10,000 events.

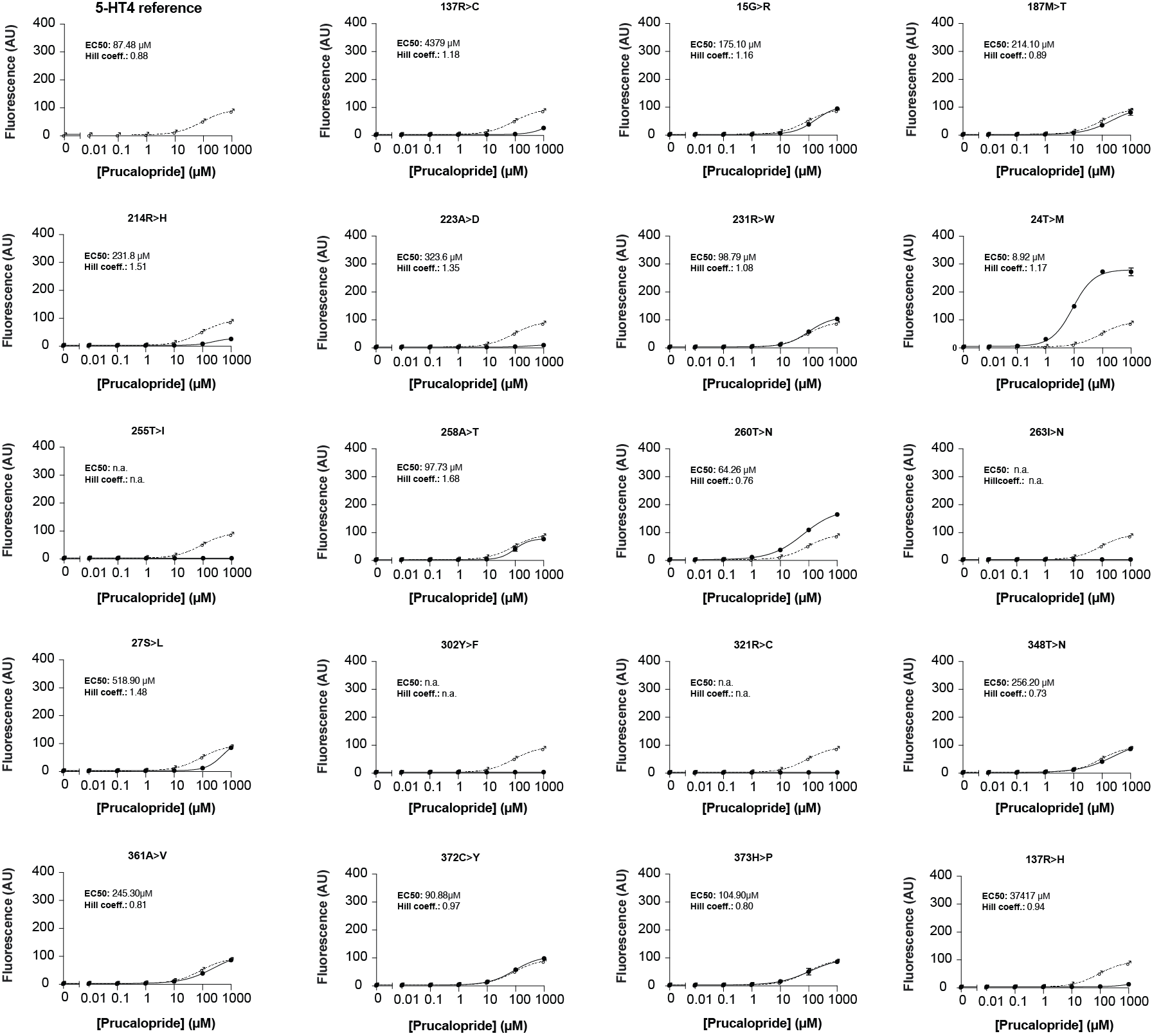

**Figure S3.** **Characterization of 5-HT4 serotonin GPCR variants from human genomes with prucalopride. A)** Dose responses with prucalopride for the 5-HT4b isoform reference receptor and 19 variants in GPA1 background. Yeast strains expressing single-amino acid variant GPCRs were induced with 0.01 μM - 1,000 μM prucalopride in addition to a non-induced “0” control. sfGFP was measured following 4 hrs of incubation with prucalopride. All data points represent median fluorescence intensity of three technical replicates (10,000 events each), from which the mean +/- the standard deviation was calculated. AU = arbitrary units, n.a. = not applicable. Data was fitted to a variable slope four-parameter curve fitting model, from which EC_50_ and Hill coefficients values were calculated. Dose-response curves show the tested variant (solid line) and the reference receptor (dashed line). The EC50 for the reference strain was 87.48 μM and the hillslope was .98.

**Supplementary Table S1**; Ensembl transcript IDs.

| **Transcript ID** | **Receptor** |
| --- | --- |
| ENST00000323865 | 5HT1A |
| ENST00000369947 | 5HT1B |
| ENST00000374619 | 5HT1D |
| ENST00000305344 | 5HT1E |
| ENST00000319595 | 5HT1F |
| ENST00000378688 | 5HT2A |
| ENST00000258400 | 5HT2B |
| ENST00000276198 | 5HT2C |
| ENST00000377888 | 5HT4 |
| ENST00000287907 | 5HT5A |
| ENST00000289753 | 5HT6 |
| ENST00000336152 | 5HT7 |

**Supplementary Table S2. Data corresponding to heatmap Figure 1C.** Mean fluorescence intensity of triplicates at both 0 and 100 μM, and the corresponding fold-change for 144 serotonin biosensor strains as explained by Gα background and plasmid.

| **Gα background** | **Plasmid** | **pH** | **Mean fluorescence intensity of triplicates**  **at 0 μM** | **Mean fluorescence intensity of triplicates**  **at 100 μM** | **Fold-change** |
| --- | --- | --- | --- | --- | --- |
| GPA1 | 5-HT1A | 4.8 | 2.72 | 2.58 | 0.95 |
| GPA1 | 5-HT1A | 7.2 | 4.18 | 4.82 | 1.15 |
| Gαs/olf | 5-HT1A | 4.8 | 2.70 | 2.59 | 0.96 |
| Gαs/olf | 5-HT1A | 7.2 | 4.01 | 4.46 | 1.11 |
| Gα12 | 5-HT1A | 4.8 | 2.59 | 2.64 | 1.02 |
| Gα12 | 5-HT1A | 7.2 | 4.10 | 4.43 | 1.08 |
| Gα13 | 5-HT1A | 4.8 | 2.74 | 2.77 | 1.01 |
| Gα13 | 5-HT1A | 7.2 | 4.19 | 4.31 | 1.03 |
| Gαi1/2 | 5-HT1A | 4.8 | 2.64 | 2.60 | 0.99 |
| Gαi1/2 | 5-HT1A | 7.2 | 4.16 | 4.48 | 1.08 |
| Gαi3 | 5-HT1A | 4.8 | 2.62 | 2.58 | 0.98 |
| Gαi3 | 5-HT1A | 7.2 | 4.21 | 4.60 | 1.09 |
| Gαz | 5-HT1A | 4.8 | 2.98 | 2.93 | 0.98 |
| Gαz | 5-HT1A | 7.2 | 4.13 | 6.19 | 1.50 |
| Gα15/16 | 5-HT1A | 4.8 | 2.63 | 2.67 | 1.01 |
| Gα15/16 | 5-HT1A | 7.2 | 3.92 | 4.49 | 1.14 |
| Gαq/11 | 5-HT1A | 4.8 | 2.66 | 2.74 | 1.03 |
| Gαq/11 | 5-HT1A | 7.2 | 4.03 | 4.15 | 1.03 |
| Gα14 | 5-HT1A | 4.8 | 2.71 | 2.67 | 0.98 |
| Gα14 | 5-HT1A | 7.2 | 4.19 | 4.69 | 1.12 |
| Gαo | 5-HT1A | 4.8 | 2.68 | 2.67 | 0.99 |
| Gαo | 5-HT1A | 7.2 | 4.12 | 4.79 | 1.16 |
| tGPA1 | 5-HT1A | 4.8 | 3.38 | 3.42 | 1.01 |
| tGPA1 | 5-HT1A | 7.2 | 4.43 | 5.13 | 1.16 |
| GPA1 | 5-HT1B | 4.8 | 2.43 | 2.39 | 0.98 |
| GPA1 | 5-HT1B | 7.2 | 5.31 | 4.57 | 0.86 |
| Gαs/olf | 5-HT1B | 4.8 | 2.41 | 2.39 | 0.99 |
| Gαs/olf | 5-HT1B | 7.2 | 3.95 | 4.34 | 1.10 |
| Gα12 | 5-HT1B | 4.8 | 2.43 | 2.37 | 0.98 |
| Gα12 | 5-HT1B | 7.2 | 3.93 | 4.02 | 1.02 |
| Gα13 | 5-HT1B | 4.8 | 2.41 | 2.39 | 0.99 |
| Gα13 | 5-HT1B | 7.2 | 3.74 | 4.08 | 1.09 |
| Gαi1/2 | 5-HT1B | 4.8 | 2.38 | 2.36 | 0.99 |
| Gαi1/2 | 5-HT1B | 7.2 | 3.54 | 4.59 | 1.30 |
| Gαi3 | 5-HT1B | 4.8 | 2.32 | 2.35 | 1.01 |
| Gαi3 | 5-HT1B | 7.2 | 3.93 | 5.81 | 1.48 |
| Gαz | 5-HT1B | 4.8 | 2.54 | 2.54 | 1.00 |
| Gαz | 5-HT1B | 7.2 | 4.81 | 5.44 | 1.13 |
| Gα15/16 | 5-HT1B | 4.8 | 2.37 | 2.28 | 0.96 |
| Gα15/16 | 5-HT1B | 7.2 | 4.49 | 4.14 | 0.92 |
| Gαq/11 | 5-HT1B | 4.8 | 2.33 | 2.28 | 0.98 |
| Gαq/11 | 5-HT1B | 7.2 | 4.04 | 4.10 | 1.02 |
| Gα14 | 5-HT1B | 4.8 | 2.45 | 2.41 | 0.98 |
| Gα14 | 5-HT1B | 7.2 | 4.08 | 3.92 | 0.96 |
| Gαo | 5-HT1B | 4.8 | 2.40 | 2.32 | 0.97 |
| Gαo | 5-HT1B | 7.2 | 4.13 | 3.95 | 0.96 |
| tGPA1 | 5-HT1B | 4.8 | 2.88 | 2.92 | 1.02 |
| tGPA1 | 5-HT1B | 7.2 | 5.31 | 4.62 | 0.87 |
| GPA1 | 5-HT1D | 4.8 | 3.58 | 3.85 | 1.08 |
| GPA1 | 5-HT1D | 7.2 | 4.82 | 4.51 | 0.93 |
| Gαs/olf | 5-HT1D | 4.8 | 3.43 | 3.45 | 1.01 |
| Gαs/olf | 5-HT1D | 7.2 | 4.13 | 4.01 | 0.97 |
| Gα12 | 5-HT1D | 4.8 | 3.64 | 4.10 | 1.13 |
| Gα12 | 5-HT1D | 7.2 | 4.25 | 4.12 | 0.97 |
| Gα13 | 5-HT1D | 4.8 | 3.63 | 3.89 | 1.07 |
| Gα13 | 5-HT1D | 7.2 | 4.23 | 4.19 | 0.99 |
| Gαi1/2 | 5-HT1D | 4.8 | 3.71 | 3.70 | 1.00 |
| Gαi1/2 | 5-HT1D | 7.2 | 3.98 | 3.70 | 0.93 |
| Gαi3 | 5-HT1D | 4.8 | 3.59 | 3.45 | 0.96 |
| Gαi3 | 5-HT1D | 7.2 | 4.06 | 3.76 | 0.93 |
| Gαz | 5-HT1D | 4.8 | 3.65 | 3.68 | 1.01 |
| Gαz | 5-HT1D | 7.2 | 4.35 | 4.53 | 1.04 |
| Gα15/16 | 5-HT1D | 4.8 | 3.42 | 3.35 | 0.98 |
| Gα15/16 | 5-HT1D | 7.2 | 3.94 | 3.64 | 0.93 |
| Gαq/11 | 5-HT1D | 4.8 | 3.54 | 3.47 | 0.98 |
| Gαq/11 | 5-HT1D | 7.2 | 4.15 | 4.27 | 1.03 |
| Gα14 | 5-HT1D | 4.8 | 3.52 | 3.48 | 0.99 |
| Gα14 | 5-HT1D | 7.2 | 4.35 | 5.05 | 1.16 |
| Gαo | 5-HT1D | 4.8 | 3.47 | 3.42 | 0.98 |
| Gαo | 5-HT1D | 7.2 | 4.49 | 4.67 | 1.04 |
| tGPA1 | 5-HT1D | 4.8 | 3.68 | 3.62 | 0.98 |
| tGPA1 | 5-HT1D | 7.2 | 4.79 | 4.90 | 1.02 |
| GPA1 | 5-HT1E | 4.8 | 3.19 | 3.07 | 0.96 |
| GPA1 | 5-HT1E | 7.2 | 4.87 | 4.79 | 0.98 |
| Gαs/olf | 5-HT1E | 4.8 | 3.09 | 2.99 | 0.97 |
| Gαs/olf | 5-HT1E | 7.2 | 4.68 | 4.82 | 1.03 |
| Gα12 | 5-HT1E | 4.8 | 3.19 | 3.14 | 0.98 |
| Gα12 | 5-HT1E | 7.2 | 4.73 | 4.14 | 0.88 |
| Gα13 | 5-HT1E | 4.8 | 3.14 | 3.10 | 0.99 |
| Gα13 | 5-HT1E | 7.2 | 4.36 | 4.37 | 1.00 |
| Gαi1/2 | 5-HT1E | 4.8 | 3.21 | 3.17 | 0.99 |
| Gαi1/2 | 5-HT1E | 7.2 | 4.22 | 5.33 | 1.26 |
| Gαi3 | 5-HT1E | 4.8 | 3.18 | 3.21 | 1.01 |
| Gαi3 | 5-HT1E | 7.2 | 4.30 | 5.01 | 1.16 |
| Gαz | 5-HT1E | 4.8 | 3.17 | 3.39 | 1.07 |
| Gαz | 5-HT1E | 7.2 | 4.69 | 7.89 | 1.68 |
| Gα15/16 | 5-HT1E | 4.8 | 3.02 | 2.96 | 0.98 |
| Gα15/16 | 5-HT1E | 7.2 | 4.77 | 4.29 | 0.90 |
| Gαq/11 | 5-HT1E | 4.8 | 3.17 | 3.13 | 0.99 |
| Gαq/11 | 5-HT1E | 7.2 | 4.80 | 4.38 | 0.91 |
| Gα14 | 5-HT1E | 4.8 | 3.23 | 3.10 | 0.96 |
| Gα14 | 5-HT1E | 7.2 | 4.83 | 4.54 | 0.94 |
| Gαo | 5-HT1E | 4.8 | 3.09 | 3.06 | 0.99 |
| Gαo | 5-HT1E | 7.2 | 4.76 | 4.71 | 0.99 |
| tGPA1 | 5-HT1E | 4.8 | 3.38 | 3.41 | 1.01 |
| tGPA1 | 5-HT1E | 7.2 | 5.43 | 5.39 | 0.99 |
| GPA1 | 5-HT1F | 4.8 | 2.88 | 2.84 | 0.99 |
| GPA1 | 5-HT1F | 7.2 | 3.93 | 3.64 | 0.92 |
| Gαs/olf | 5-HT1F | 4.8 | 2.97 | 2.99 | 1.01 |
| Gαs/olf | 5-HT1F | 7.2 | 3.47 | 3.55 | 1.02 |
| Gα12 | 5-HT1F | 4.8 | 3.00 | 2.96 | 0.99 |
| Gα12 | 5-HT1F | 7.2 | 3.52 | 3.65 | 1.04 |
| Gα13 | 5-HT1F | 4.8 | 3.10 | 3.04 | 0.98 |
| Gα13 | 5-HT1F | 7.2 | 3.98 | 3.68 | 0.92 |
| Gαi1/2 | 5-HT1F | 4.8 | 2.78 | 2.83 | 1.02 |
| Gαi1/2 | 5-HT1F | 7.2 | 3.40 | 3.58 | 1.05 |
| Gαi3 | 5-HT1F | 4.8 | 2.76 | 2.81 | 1.02 |
| Gαi3 | 5-HT1F | 7.2 | 3.49 | 3.15 | 0.90 |
| Gαz | 5-HT1F | 4.8 | 3.05 | 2.97 | 0.97 |
| Gαz | 5-HT1F | 7.2 | 4.08 | 3.75 | 0.92 |
| Gα15/16 | 5-HT1F | 4.8 | 2.80 | 2.74 | 0.98 |
| Gα15/16 | 5-HT1F | 7.2 | 3.26 | 3.07 | 0.94 |
| Gαq/11 | 5-HT1F | 4.8 | 2.99 | 2.97 | 0.99 |
| Gαq/11 | 5-HT1F | 7.2 | 3.28 | 3.18 | 0.97 |
| Gα14 | 5-HT1F | 4.8 | 3.05 | 3.06 | 1.00 |
| Gα14 | 5-HT1F | 7.2 | 3.70 | 3.57 | 0.97 |
| Gαo | 5-HT1F | 4.8 | 2.87 | 2.83 | 0.99 |
| Gαo | 5-HT1F | 7.2 | 3.37 | 3.55 | 1.05 |
| tGPA1 | 5-HT1F | 4.8 | 3.23 | 3.21 | 0.99 |
| tGPA1 | 5-HT1F | 7.2 | 4.75 | 4.35 | 0.92 |
| GPA1 | 5-HT2A | 4.8 | 2.49 | 2.38 | 0.96 |
| GPA1 | 5-HT2A | 7.2 | 4.35 | 4.27 | 0.98 |
| Gαs/olf | 5-HT2A | 4.8 | 2.38 | 2.34 | 0.98 |
| Gαs/olf | 5-HT2A | 7.2 | 4.38 | 4.53 | 1.03 |
| Gα12 | 5-HT2A | 4.8 | 2.47 | 2.41 | 0.98 |
| Gα12 | 5-HT2A | 7.2 | 3.91 | 3.56 | 0.91 |
| Gα13 | 5-HT2A | 4.8 | 2.52 | 2.46 | 0.97 |
| Gα13 | 5-HT2A | 7.2 | 4.11 | 3.53 | 0.86 |
| Gαi1/2 | 5-HT2A | 4.8 | 2.41 | 2.36 | 0.98 |
| Gαi1/2 | 5-HT2A | 7.2 | 4.06 | 3.82 | 0.94 |
| Gαi3 | 5-HT2A | 4.8 | 2.42 | 2.36 | 0.98 |
| Gαi3 | 5-HT2A | 7.2 | 5.19 | 5.12 | 0.99 |
| Gαz | 5-HT2A | 4.8 | 2.55 | 2.50 | 0.98 |
| Gαz | 5-HT2A | 7.2 | 4.35 | 5.03 | 1.16 |
| Gα15/16 | 5-HT2A | 4.8 | 2.35 | 2.28 | 0.97 |
| Gα15/16 | 5-HT2A | 7.2 | 3.73 | 4.20 | 1.13 |
| Gαq/11 | 5-HT2A | 4.8 | 2.45 | 2.40 | 0.98 |
| Gαq/11 | 5-HT2A | 7.2 | 4.21 | 4.93 | 1.17 |
| Gα14 | 5-HT2A | 4.8 | 2.42 | 2.36 | 0.98 |
| Gα14 | 5-HT2A | 7.2 | 4.62 | 5.33 | 1.16 |
| Gαo | 5-HT2A | 4.8 | 2.43 | 2.38 | 0.98 |
| Gαo | 5-HT2A | 7.2 | 3.61 | 4.96 | 1.37 |
| tGPA1 | 5-HT2A | 4.8 | 3.05 | 2.93 | 0.96 |
| tGPA1 | 5-HT2A | 7.2 | 3.87 | 4.22 | 1.09 |
| GPA1 | 5-HT2B | 4.8 | 2.64 | 2.55 | 0.96 |
| GPA1 | 5-HT2B | 7.2 | 4.30 | 4.24 | 0.99 |
| Gαs/olf | 5-HT2B | 4.8 | 2.60 | 2.58 | 0.99 |
| Gαs/olf | 5-HT2B | 7.2 | 3.82 | 4.05 | 1.06 |
| Gα12 | 5-HT2B | 4.8 | 2.54 | 2.50 | 0.98 |
| Gα12 | 5-HT2B | 7.2 | 3.85 | 4.17 | 1.08 |
| Gα13 | 5-HT2B | 4.8 | 2.59 | 2.63 | 1.02 |
| Gα13 | 5-HT2B | 7.2 | 4.07 | 3.99 | 0.98 |
| Gαi1/2 | 5-HT2B | 4.8 | 2.72 | 2.48 | 0.91 |
| Gαi1/2 | 5-HT2B | 7.2 | 3.82 | 3.78 | 0.99 |
| Gαi3 | 5-HT2B | 4.8 | 2.56 | 2.42 | 0.95 |
| Gαi3 | 5-HT2B | 7.2 | 3.75 | 3.75 | 1.00 |
| Gαz | 5-HT2B | 4.8 | 3.04 | 2.74 | 0.90 |
| Gαz | 5-HT2B | 7.2 | 4.33 | 4.40 | 1.02 |
| Gα15/16 | 5-HT2B | 4.8 | 2.51 | 2.30 | 0.92 |
| Gα15/16 | 5-HT2B | 7.2 | 3.63 | 3.55 | 0.98 |
| Gαq/11 | 5-HT2B | 4.8 | 2.60 | 2.57 | 0.99 |
| Gαq/11 | 5-HT2B | 7.2 | 3.79 | 3.90 | 1.03 |
| Gα14 | 5-HT2B | 4.8 | 2.69 | 2.50 | 0.93 |
| Gα14 | 5-HT2B | 7.2 | 3.88 | 3.91 | 1.01 |
| Gαo | 5-HT2B | 4.8 | 2.69 | 2.38 | 0.88 |
| Gαo | 5-HT2B | 7.2 | 3.84 | 3.91 | 1.02 |
| tGPA1 | 5-HT2B | 4.8 | 3.28 | 2.99 | 0.91 |
| tGPA1 | 5-HT2B | 7.2 | 3.94 | 3.90 | 0.99 |
| GPA1 | 5-HT2C | 4.8 | 2.89 | 2.72 | 0.94 |
| GPA1 | 5-HT2C | 7.2 | 3.55 | 4.81 | 1.36 |
| Gαs/olf | 5-HT2C | 4.8 | 2.79 | 2.84 | 1.02 |
| Gαs/olf | 5-HT2C | 7.2 | 4.12 | 4.53 | 1.10 |
| Gα12 | 5-HT2C | 4.8 | 2.83 | 2.70 | 0.95 |
| Gα12 | 5-HT2C | 7.2 | 4.21 | 3.79 | 0.90 |
| Gα13 | 5-HT2C | 4.8 | 3.05 | 3.04 | 1.00 |
| Gα13 | 5-HT2C | 7.2 | 4.10 | 3.98 | 0.97 |
| Gαi1/2 | 5-HT2C | 4.8 | 2.98 | 2.86 | 0.96 |
| Gαi1/2 | 5-HT2C | 7.2 | 3.74 | 4.32 | 1.16 |
| Gαi3 | 5-HT2C | 4.8 | 2.96 | 2.88 | 0.97 |
| Gαi3 | 5-HT2C | 7.2 | 3.68 | 3.31 | 0.90 |
| Gαz | 5-HT2C | 4.8 | 3.40 | 3.44 | 1.01 |
| Gαz | 5-HT2C | 7.2 | 4.12 | 3.69 | 0.90 |
| Gα15/16 | 5-HT2C | 4.8 | 2.79 | 2.86 | 1.02 |
| Gα15/16 | 5-HT2C | 7.2 | 3.38 | 3.12 | 0.93 |
| Gαq/11 | 5-HT2C | 4.8 | 3.30 | 3.33 | 1.01 |
| Gαq/11 | 5-HT2C | 7.2 | 3.53 | 3.37 | 0.96 |
| Gα14 | 5-HT2C | 4.8 | 3.07 | 2.98 | 0.97 |
| Gα14 | 5-HT2C | 7.2 | 3.60 | 3.62 | 1.01 |
| Gαo | 5-HT2C | 4.8 | 2.75 | 2.68 | 0.97 |
| Gαo | 5-HT2C | 7.2 | 4.11 | 3.89 | 0.95 |
| tGPA1 | 5-HT2C | 4.8 | 3.15 | 3.04 | 0.96 |
| tGPA1 | 5-HT2C | 7.2 | 4.12 | 3.49 | 0.85 |
| GPA1 | 5-HT4 | 4.8 | 3.10 | 141.63 | 45.62 |
| GPA1 | 5-HT4 | 7.2 | 11.15 | 143.08 | 12.83 |
| Gαs/olf | 5-HT4 | 4.8 | 3.26 | 5.92 | 1.81 |
| Gαs/olf | 5-HT4 | 7.2 | 4.92 | 105.25 | 21.41 |
| Gα12 | 5-HT4 | 4.8 | 3.36 | 13.38 | 3.98 |
| Gα12 | 5-HT4 | 7.2 | 4.91 | 117.65 | 23.95 |
| Gα13 | 5-HT4 | 4.8 | 3.07 | 24.55 | 7.99 |
| Gα13 | 5-HT4 | 7.2 | 4.91 | 161.90 | 32.97 |
| Gαi1/2 | 5-HT4 | 4.8 | 3.40 | 142.22 | 41.84 |
| Gαi1/2 | 5-HT4 | 7.2 | 18.59 | 286.34 | 15.40 |
| Gαi3 | 5-HT4 | 4.8 | 3.29 | 158.46 | 48.15 |
| Gαi3 | 5-HT4 | 7.2 | 5.80 | 97.56 | 16.83 |
| Gαz | 5-HT4 | 4.8 | 3.33 | 212.57 | 63.84 |
| Gαz | 5-HT4 | 7.2 | 72.43 | 291.58 | 4.03 |
| Gα15/16 | 5-HT4 | 4.8 | 3.20 | 88.70 | 27.75 |
| Gα15/16 | 5-HT4 | 7.2 | 7.67 | 42.60 | 5.56 |
| Gαq/11 | 5-HT4 | 4.8 | 3.30 | 8.93 | 2.71 |
| Gαq/11 | 5-HT4 | 7.2 | 4.29 | 112.63 | 26.28 |
| Gα14 | 5-HT4 | 4.8 | 3.03 | 25.46 | 8.40 |
| Gα14 | 5-HT4 | 7.2 | 4.56 | 231.00 | 50.68 |
| Gαo | 5-HT4 | 4.8 | 3.24 | 18.84 | 5.81 |
| Gαo | 5-HT4 | 7.2 | 4.88 | 200.01 | 41.00 |
| tGPA1 | 5-HT4 | 4.8 | 3.74 | 3.64 | 0.97 |
| tGPA1 | 5-HT4 | 7.2 | 5.56 | 5.58 | 1.00 |
| GPA1 | 5-HT5A | 4.8 | 2.85 | 2.71 | 0.95 |
| GPA1 | 5-HT5A | 7.2 | 4.82 | 4.10 | 0.85 |
| Gαs/olf | 5-HT5A | 4.8 | 2.82 | 2.86 | 1.01 |
| Gαs/olf | 5-HT5A | 7.2 | 4.04 | 4.51 | 1.12 |
| Gα12 | 5-HT5A | 4.8 | 2.76 | 2.68 | 0.97 |
| Gα12 | 5-HT5A | 7.2 | 3.95 | 4.79 | 1.21 |
| Gα13 | 5-HT5A | 4.8 | 2.83 | 2.81 | 0.99 |
| Gα13 | 5-HT5A | 7.2 | 4.36 | 4.64 | 1.07 |
| Gαi1/2 | 5-HT5A | 4.8 | 2.75 | 2.74 | 1.00 |
| Gαi1/2 | 5-HT5A | 7.2 | 4.17 | 4.52 | 1.08 |
| Gαi3 | 5-HT5A | 4.8 | 2.87 | 2.87 | 1.00 |
| Gαi3 | 5-HT5A | 7.2 | 4.05 | 4.26 | 1.05 |
| Gαz | 5-HT5A | 4.8 | 3.45 | 3.55 | 1.03 |
| Gαz | 5-HT5A | 7.2 | 4.54 | 4.95 | 1.09 |
| Gα15/16 | 5-HT5A | 4.8 | 2.75 | 2.73 | 0.99 |
| Gα15/16 | 5-HT5A | 7.2 | 3.43 | 3.91 | 1.14 |
| Gαq/11 | 5-HT5A | 4.8 | 2.80 | 2.80 | 1.00 |
| Gαq/11 | 5-HT5A | 7.2 | 3.73 | 3.69 | 0.99 |
| Gα14 | 5-HT5A | 4.8 | 2.89 | 2.87 | 0.99 |
| Gα14 | 5-HT5A | 7.2 | 3.89 | 3.75 | 0.96 |
| Gαo | 5-HT5A | 4.8 | 2.79 | 2.69 | 0.96 |
| Gαo | 5-HT5A | 7.2 | 3.94 | 3.92 | 0.99 |
| tGPA1 | 5-HT5A | 4.8 | 3.79 | 3.82 | 1.01 |
| tGPA1 | 5-HT5A | 7.2 | 3.85 | 4.75 | 1.24 |
| GPA1 | 5-HT6 | 4.8 | 2.81 | 2.84 | 1.01 |
| GPA1 | 5-HT6 | 7.2 | 4.16 | 4.19 | 1.01 |
| Gαs/olf | 5-HT6 | 4.8 | 3.05 | 3.05 | 1.00 |
| Gαs/olf | 5-HT6 | 7.2 | 4.51 | 4.67 | 1.03 |
| Gα12 | 5-HT6 | 4.8 | 2.79 | 2.80 | 1.00 |
| Gα12 | 5-HT6 | 7.2 | 4.75 | 4.46 | 0.94 |
| Gα13 | 5-HT6 | 4.8 | 3.49 | 3.35 | 0.96 |
| Gα13 | 5-HT6 | 7.2 | 4.30 | 4.42 | 1.03 |
| Gαi1/2 | 5-HT6 | 4.8 | 2.88 | 2.83 | 0.98 |
| Gαi1/2 | 5-HT6 | 7.2 | 3.96 | 3.86 | 0.98 |
| Gαi3 | 5-HT6 | 4.8 | 2.82 | 2.85 | 1.01 |
| Gαi3 | 5-HT6 | 7.2 | 4.32 | 3.67 | 0.85 |
| Gαz | 5-HT6 | 4.8 | 3.28 | 3.21 | 0.98 |
| Gαz | 5-HT6 | 7.2 | 4.57 | 4.14 | 0.91 |
| Gα15/16 | 5-HT6 | 4.8 | 2.77 | 2.77 | 1.00 |
| Gα15/16 | 5-HT6 | 7.2 | 3.69 | 3.36 | 0.91 |
| Gαq/11 | 5-HT6 | 4.8 | 2.83 | 2.86 | 1.01 |
| Gαq/11 | 5-HT6 | 7.2 | 3.96 | 3.11 | 0.78 |
| Gα14 | 5-HT6 | 4.8 | 2.97 | 2.97 | 1.00 |
| Gα14 | 5-HT6 | 7.2 | 4.36 | 4.03 | 0.92 |
| Gαo | 5-HT6 | 4.8 | 2.85 | 2.86 | 1.00 |
| Gαo | 5-HT6 | 7.2 | 4.51 | 4.49 | 0.99 |
| tGPA1 | 5-HT6 | 4.8 | 3.44 | 3.67 | 1.07 |
| tGPA1 | 5-HT6 | 7.2 | 4.92 | 5.11 | 1.04 |
| GPA1 | 5-HT7 | 4.8 | 2.90 | 2.97 | 1.02 |
| GPA1 | 5-HT7 | 7.2 | 4.68 | 5.40 | 1.15 |
| Gαs/olf | 5-HT7 | 4.8 | 3.52 | 3.45 | 0.98 |
| Gαs/olf | 5-HT7 | 7.2 | 5.04 | 4.11 | 0.82 |
| Gα12 | 5-HT7 | 4.8 | 2.88 | 2.84 | 0.99 |
| Gα12 | 5-HT7 | 7.2 | 4.04 | 4.86 | 1.20 |
| Gα13 | 5-HT7 | 4.8 | 2.94 | 2.84 | 0.97 |
| Gα13 | 5-HT7 | 7.2 | 4.49 | 4.53 | 1.01 |
| Gαi1/2 | 5-HT7 | 4.8 | 2.89 | 2.85 | 0.98 |
| Gαi1/2 | 5-HT7 | 7.2 | 4.05 | 4.89 | 1.21 |
| Gαi3 | 5-HT7 | 4.8 | 2.93 | 2.74 | 0.94 |
| Gαi3 | 5-HT7 | 7.2 | 4.55 | 4.64 | 1.02 |
| Gαz | 5-HT7 | 4.8 | 3.64 | 3.53 | 0.97 |
| Gαz | 5-HT7 | 7.2 | 4.79 | 4.78 | 1.00 |
| Gα15/16 | 5-HT7 | 4.8 | 2.86 | 2.80 | 0.98 |
| Gα15/16 | 5-HT7 | 7.2 | 4.54 | 4.06 | 0.89 |
| Gαq/11 | 5-HT7 | 4.8 | 2.95 | 3.00 | 1.02 |
| Gαq/11 | 5-HT7 | 7.2 | 4.02 | 4.13 | 1.03 |
| Gα14 | 5-HT7 | 4.8 | 3.12 | 3.11 | 1.00 |
| Gα14 | 5-HT7 | 7.2 | 4.09 | 3.92 | 0.96 |
| Gαo | 5-HT7 | 4.8 | 2.96 | 2.97 | 1.01 |
| Gαo | 5-HT7 | 7.2 | 4.27 | 4.74 | 1.11 |
| tGPA1 | 5-HT7 | 4.8 | 3.72 | 3.57 | 0.96 |
| tGPA1 | 5-HT7 | 7.2 | 5.17 | 5.76 | 1.11 |

**Supplementary Table S3. EC_50_s and *R^2^* for pH data**

| **Strain + pH** | **pH** | **R^2^** | **EC_50_** |
| --- | --- | --- | --- |
| 5-HT4+Gαz (=yBL167) | 2 | 0.02 | Unstable |
| 5-HT4+Gαz (=yBL167) | 3 | 0.98 | 46.99 |
| 5-HT4+Gαz (=yBL167) | 4 | 0.99 | 11.17 |
| 5-HT4+Gαz (=yBL167) | 5 | 0.99 | 3.87 |
| 5-HT4+Gαz (=yBL167) | 6 | 0.95 | 0.60 |
| 5-HT4+Gαz (=yBL167) | 7 | 0.90 | 0.01 |

**Supplementary Table S4: R^2,^ EC_50_, and Hill coeff. from spent medium test**

| **Condition** | **R^2^** | **EC_50_** | **Hillslope** |
| --- | --- | --- | --- |
| 10% spiked MQ | 0.9994 | 4.0010 | 1.0870 |
| 10% spiked 72h SM | 0.9957 | 7.8790 | 1.4990 |
| 25% spiked MQ | 0.9944 | 2.5940 | 1.1120 |
| 25% spiked 72h SM | 0.9997 | 11.2100 | 1.4110 |
| 50% spiked MQ | 0.9962 | 1.6970 | 1.0230 |
| 50% spiked 72h SM | 0.9995 | 20.4100 | 1.1970 |

**Supplementary Table S5: gBlocks for serotonin GPCRs, based on UniProt entries, including DNA sequence.**

| **gBlock number** | **Name** | **Based on UniprotID** | **DNA sequence** |
| --- | --- | --- | --- |
| gBL007 | 5-hydroxytryptamine receptor 1A | P08908-1 (canonical) | AAAACAATGGACGTATTATCACCTGGACAGGGCAACAACACAACAAGTCCCCCTGCACCTTTCGAGACAGGCGGGAACACAACAGGCATCAGTGACGTGACAGTATCATACCAGGTGATCACATCCTTACTACTTGGGACATTAATATTCTGCGCCGTCCTGGGAAACGCATGCGTAGTGGCAGCCATAGCCCTGGAGAGGAGTCTGCAGAACGTCGCAAACTACCTGATCGGCAGCTTAGCCGTAACAGACTTAATGGTCTCAGTGTTAGTGCTTCCGATGGCCGCATTATACCAGGTGCTAAACAAGTGGACGCTGGGACAGGTGACATGCGACCTTTTCATCGCGTTAGACGTCTTATGCTGCACATCATCGATCCTTCACCTTTGCGCAATCGCCCTTGACCGTTACTGGGCCATCACCGACCCCATCGACTACGTGAACAAGAGGACCCCTAGGCGTGCGGCCGCATTAATATCGCTGACCTGGTTAATCGGATTCCTAATATCCATACCGCCTATGTTAGGATGGAGAACCCCGGAAGACCGTAGCGACCCCGACGCCTGCACGATAAGCAAGGACCACGGATACACCATCTACAGTACATTCGGAGCATTCTACATCCCTTTATTACTAATGTTAGTACTTTACGGCCGTATATTCCGTGCCGCAAGGTTCAGGATCCGTAAGACAGTAAAGAAGGTCGAGAAGACCGGAGCAGACACAAGACACGGGGCCAGTCCTGCTCCGCAGCCAAAGAAGAGTGTGAACGGCGAGAGTGGGTCGAGGAATTGGAGACTGGGCGTCGAGTCGAAGGCCGGCGGAGCATTATGTGCTAACGGGGCGGTCAGGCAAGGTGACGACGGCGCAGCGTTAGAGGTAATAGAGGTCCACAGGGTCGGAAACAGCAAGGAGCACTTACCTTTACCTTCTGAGGCCGGACCTACCCCGTGCGCGCCTGCCAGCTTCGAGAGGAAGAACGAGAGGAACGCAGAGGCGAAGCGTAAGATGGCCTTAGCAAGGGAGCGTAAGACAGTGAAGACACTAGGCATCATAATGGGAACCTTCATACTATGCTGGTTACCTTTCTTCATAGTCGCCCTTGTCCTTCCCTTCTGCGAGTCGTCCTGCCACATGCCGACATTACTTGGGGCAATCATCAACTGGCTTGGCTACAGTAACTCCCTTCTAAACCCCGTGATCTACGCCTACTTCAACAAGGACTTCCAGAACGCCTTCAAGAAGATAATCAAGTGCAAGTTCTGCCGTCAGTAG |
| gBL021 | 5-hydroxytryptamine receptor 1B | P28222-1 (canonical) | AAAACAATGGAAGAACCAGGAGCCCAATGTGCGCCACCACCACCAGCAGGTTCCGAAACTTGGGTTCCGCAAGCCAATCTGTCTTCTGCTCCATCCCAAAATTGCTCTGCTAAAGATTATATATATCAAGATTCCATTTCACTGCCATGGAAAGTTTTGTTGGTTATGTTGTTAGCGCTGATTACTTTAGCTACCACGCTTTCCAATGCTTTTGTTATTGCAACTGTATATAGAACTAGAAAATTGCACACGCCCGCGAATTATCTGATTGCTTCTTTAGCCGTCACCGATTTGCTGGTCTCAATTTTGGTTATGCCTATTTCTACCATGTATACTGTTACCGGCAGATGGACTCTTGGTCAAGTGGTTTGTGACTTTTGGCTTAGTAGCGATATTACTTGTTGTACGGCCTCAATTCTGCATTTATGTGTTATTGCACTAGATCGTTATTGGGCTATTACCGATGCCGTTGAATATTCGGCGAAAAGAACTCCTAAACGTGCTGCAGTAATGATTGCCCTAGTATGGGTATTTTCTATTTCGATTTCCTTGCCGCCGTTCTTTTGGAGACAAGCCAAAGCAGAGGAAGAAGTAAGCGAATGTGTTGTCAATACTGATCATATCCTTTATACTGTGTATTCAACTGTCGGAGCCTTCTATTTTCCTACCCTTTTACTGATTGCACTATATGGTAGAATTTATGTCGAGGCTAGATCAAGAATTCTTAAACAAACGCCAAATAGAACAGGCAAAAGATTGACTCGTGCACAATTGATTACGGATAGCCCAGGCAGCACCTCTTCTGTTACAAGTATTAATTCTAGAGTTCCCGATGTACCGTCCGAATCCGGCTCACCAGTTTATGTCAATCAAGTTAAAGTTAGAGTATCTGATGCCTTGCTTGAAAAGAAGAAATTGATGGCAGCCAGAGAAAGAAAAGCTACGAAAACCTTGGGGATTATCCTAGGTGCTTTTATTGTATGTTGGCTACCATTCTTTATTATCTCTCTGGTTATGCCAATTTGTAAAGATGCCTGTTGGTTTCATTTGGCAATCTTTGACTTCTTCACGTGGTTAGGGTATCTGAATTCACTAATAAACCCAATTATTTATACCATGTCGAATGAAGATTTTAAACAAGCCTTCCATAAACTAATAAGGTTTAAGTGTACCAGTTAG |
| gBL022 | 5-hydroxytryptamine receptor 1D | P28221-1 (canonical) | AAAACAATGTCTCCATTGAATCAATCTGCTGAAGGTTTGCCACAAGAAGCTTCTAATAGATCTTTGAATGCTACTGAAACTTCTGAAGCTTGGGATCCAAGAACTTTGCAAGCTTTGAAAATTTCTTTGGCTGTTGTTTTGTCTGTTATTACTTTGGCTACTGTTTTGTCTAATGCTTTTGTTTTGACTACTATTTTGTTGACTAGAAAATTGCATACTCCAGCTAATTATTTGATTGGTTCTTTGGCTACTACTGATTTGTTGGTTTCTATTTTGGTTATGCCAATTTCTATTGCTTATACTATTACTCATACTTGGAATTTTGGTCAAATTTTGTGTGATATTTGGTTGTCTTCTGATATTACTTGTTGTACTGCTTCTATTTTGCATTTGTGTGTTATTGCTTTGGATAGATATTGGGCTATTACTGATGCTTTGGAATATTCTAAAAGAAGAACTGCTGGTCATGCTGCTACTATGATTGCTATTGTTTGGGCTATTTCTATTTGTATTTCTATTCCACCATTGTTTTGGAGACAAGCTAAAGCTCAAGAAGAAATGTCTGATTGTTTGGTTAATACTTCTCAAATTTCTTATACTATTTATTCTACTTGTGGTGCTTTTTATATTCCATCTGTTTTGTTGATTATTTTGTATGGTAGAATTTATAGAGCTGCTAGAAATAGAATTTTGAATCCACCATCTTTGTATGGTAAAAGATTTACTACTGCTCATTTGATTACTGGTTCTGCTGGTTCTTCTTTGTGTTCTTTGAATTCTTCTTTGCATGAAGGTCATTCTCATTCTGCTGGTTCTCCATTGTTTTTTAATCATGTTAAAATTAAATTGGCTGATTCTGCTTTGGAAAGAAAAAGAATTTCTGCTGCTAGAGAAAGAAAAGCTACTAAAATTTTGGGTATTATTTTGGGTGCTTTTATTATTTGTTGGTTGCCATTTTTTGTTGTTTCTTTGGTTTTGCCAATTTGTAGAGATTCTTGTTGGATTCATCCAGCTTTGTTTGATTTTTTTACTTGGTTGGGTTATTTGAATTCTTTGATTAATCCAATTATTTATACTGTTTTTAATGAAGAATTTAGACAAGCTTTTCAAAAAATTGTTCCATTTAGAAAAGCTTCTTAA |
| gBL023 | 5-hydroxytryptamine receptor 1E | P28566-1 (canonical) | AAAACAATGAATATTACTAATTGTACTACTGAAGCTTCTATGGCTATTAGACCAAAAACTATTACTGAAAAAATGTTGATTTGTATGACTTTGGTTGTTATTACTACTTTGACTACTTTGTTGAATTTGGCTGTTATTATGGCTATTGGTACTACTAAAAAATTGCATCAACCAGCTAATTATTTGATTTGTTCTTTGGCTGTTACTGATTTGTTGGTTGCTGTTTTGGTTATGCCATTGTCTATTATTTATATTGTTATGGATAGATGGAAATTGGGTTATTTTTTGTGTGAAGTTTGGTTGTCTGTTGATATGACTTGTTGTACTTGTTCTATTTTGCATTTGTGTGTTATTGCTTTGGATAGATATTGGGCTATTACTAATGCTATTGAATATGCTAGAAAAAGAACTGCTAAAAGAGCTGCTTTGATGATTTTGACTGTTTGGACTATTTCTATTTTTATTTCTATGCCACCATTGTTTTGGAGATCTCATAGAAGATTGTCTCCACCACCATCTCAATGTACTATTCAACATGATCATGTTATTTATACTATTTATTCTACTTTGGGTGCTTTTTATATTCCATTGACTTTGATTTTGATTTTGTATTATAGAATTTATCATGCTGCTAAATCTTTGTATCAAAAAAGAGGTTCTTCTAGACATTTGTCTAATAGATCTACTGATTCTCAAAATTCTTTTGCTTCTTGTAAATTGACTCAAACTTTTTGTGTTTCTGATTTTTCTACTTCTGATCCAACTACTGAATTTGAAAAATTTCATGCTTCTATTAGAATTCCACCATTTGATAATGATTTGGATCATCCAGGTGAAAGACAACAAATTTCTTCTACTAGAGAAAGAAAAGCTGCTAGAATTTTGGGTTTGATTTTGGGTGCTTTTATTTTGTCTTGGTTGCCATTTTTTATTAAAGAATTGATTGTTGGTTTGTCTATTTATACTGTTTCTTCTGAAGTTGCTGATTTTTTGACTTGGTTGGGTTATGTTAATTCTTTGATTAATCCATTGTTGTATACTTCTTTTAATGAAGATTTTAAATTGGCTTTTAAAAAATTGATTAGATGTAGAGAACATACTTAA |
| gBL024 | 5-hydroxytryptamine receptor 1F | P30939-1 (canonical) | AAAACAATGGATTTTTTGAATTCTTCTGATCAAAATTTGACTTCTGAAGAATTGTTGAATAGAATGCCATCTAAAATTTTGGTTTCTTTGACTTTGTCTGGTTTGGCTTTGATGACTACTACTATTAATTCTTTGGTTATTGCTGCTATTATTGTTACTAGAAAATTGCATCATCCAGCTAATTATTTGATTTGTTCTTTGGCTGTTACTGATTTTTTGGTTGCTGTTTTGGTTATGCCATTTTCTATTGTTTATATTGTTAGAGAATCTTGGATTATGGGTCAAGTTGTTTGTGATATTTGGTTGTCTGTTGATATTACTTGTTGTACTTGTTCTATTTTGCATTTGTCTGCTATTGCTTTGGATAGATATAGAGCTATTACTGATGCTGTTGAATATGCTAGAAAAAGAACTCCAAAACATGCTGGTATTATGATTACTATTGTTTGGATTATTTCTGTTTTTATTTCTATGCCACCATTGTTTTGGAGACATCAAGGTACTTCTAGAGATGATGAATGTATTATTAAACATGATCATATTGTTTCTACTATTTATTCTACTTTTGGTGCTTTTTATATTCCATTGGCTTTGATTTTGATTTTGTATTATAAAATTTATAGAGCTGCTAAAACTTTGTATCATAAAAGACAAGCTTCTAGAATTGCTAAAGAAGAAGTTAATGGTCAAGTTTTGTTGGAATCTGGTGAAAAATCTACTAAATCTGTTTCTACTTCTTATGTTTTGGAAAAATCTTTGTCTGATCCATCTACTGATTTTGATAAAATTCATTCTACTGTTAGATCTTTGAGATCTGAATTTAAACATGAAAAATCTTGGAGAAGACAAAAAATTTCTGGTACTAGAGAAAGAAAAGCTGCTACTACTTTGGGTTTGATTTTGGGTGCTTTTGTTATTTGTTGGTTGCCATTTTTTGTTAAAGAATTGGTTGTTAATGTTTGTGATAAATGTAAAATTTCTGAAGAAATGTCTAATTTTTTGGCTTGGTTGGGTTATTTGAATTCTTTGATTAATCCATTGATTTATACTATTTTTAATGAAGATTTTAAAAAAGCTTTTCAAAAATTGGTTAGATGTAGATGTTAA |
| gBL025 | 5-hydroxytryptamine receptor 2A | P28223-1 (canonical) | AAAACAATGGACATATTATGCGAGGAGAACACGAGTTTAAGTTCCACAACGAACAGTCTTATGCAGTTAAACGACGACACCCGTTTATACTCCAACGACTTCAACTCCGGCGAGGCAAACACCTCAGACGCGTTCAACTGGACGGTCGACTCGGAGAACAGGACCAACCTAAGCTGCGAGGGGTGCCTGTCGCCGTCATGCCTGAGTCTACTTCACCTGCAGGAGAAGAACTGGAGTGCACTACTAACCGCAGTAGTCATAATACTGACAATCGCAGGAAACATATTAGTAATCATGGCAGTAAGTCTTGAGAAGAAGTTACAGAACGCAACAAACTACTTCTTAATGTCCCTAGCAATAGCAGACATGCTTCTGGGCTTCCTAGTAATGCCTGTCTCGATGCTGACAATCCTTTACGGGTACAGGTGGCCCTTACCCTCAAAGCTGTGCGCGGTGTGGATCTACCTAGACGTGTTATTCAGCACCGCATCGATAATGCACTTATGCGCAATAAGCTTAGACAGGTACGTAGCCATACAGAACCCCATCCACCACTCACGTTTCAACAGCAGGACAAAGGCGTTCCTTAAGATAATAGCGGTCTGGACCATATCAGTGGGAATCTCAATGCCGATCCCGGTATTCGGGTTACAGGACGACTCGAAGGTCTTCAAGGAAGGCTCCTGCCTACTGGCCGACGACAACTTCGTATTAATCGGGTCGTTCGTCTCATTCTTCATCCCGTTAACAATCATGGTAATAACCTACTTCTTAACCATAAAGAGTCTACAGAAAGAGGCCACGTTATGCGTCTCCGACTTAGGCACACGTGCGAAGTTAGCGTCATTCTCGTTCCTTCCTCAGAGCTCGCTTAGTTCAGAGAAGCTTTTCCAGCGTTCCATACACCGTGAGCCCGGATCATACACCGGCAGGCGTACAATGCAGAGTATATCGAACGAGCAGAAGGCCTGCAAGGTACTGGGAATAGTGTTCTTCTTATTCGTCGTCATGTGGTGCCCTTTCTTCATAACAAACATCATGGCCGTAATATGCAAGGAGAGTTGCAACGAGGACGTCATCGGAGCATTACTTAACGTGTTCGTCTGGATAGGGTACTTATCAAGTGCCGTGAACCCTCTGGTGTACACGCTGTTCAACAAGACATACCGTTCGGCCTTCTCACGTTACATCCAGTGCCAGTACAAGGAGAACAAGAAGCCTCTTCAGTTAATCTTAGTCAACACCATACCTGCATTAGCATACAAGAGCAGCCAGTTACAGATGGGCCAGAAGAAGAACAGTAAGCAGGACGCAAAGACGACCGACAACGACTGCAGTATGGTAGCACTGGGGAAGCAGCACTCGGAAGAGGCAAGTAAGGACAACTCCGACGGCGTGAACGAGAAGGTATCATGCGTGTGA |
| gBL026 | 5-hydroxytryptamine receptor 2B | P41595-1 (canonical) | AAAACAATGGCACTGAGTTACAGGGTGAGCGAGTTACAGTCCACAATCCCTGAGCACATCCTGCAGTCAACCTTCGTACACGTAATAAGCTCAAACTGGTCCGGGCTACAGACAGAGTCAATCCCAGAGGAGATGAAGCAGATCGTCGAGGAGCAGGGAAACAAGTTACACTGGGCAGCATTACTGATATTAATGGTCATAATACCTACAATAGGTGGAAACACCTTAGTGATACTGGCAGTGAGCTTAGAGAAGAAGCTGCAGTACGCGACCAACTACTTCTTAATGAGTTTAGCCGTCGCAGACCTATTAGTCGGGTTATTCGTAATGCCTATCGCATTACTGACAATCATGTTCGAGGCGATGTGGCCTCTTCCGTTAGTCTTATGCCCCGCATGGCTTTTCCTGGACGTACTATTCTCAACAGCCAGTATCATGCACCTATGCGCAATATCCGTAGACAGGTACATAGCCATCAAGAAGCCTATCCAGGCCAACCAGTACAACTCGAGGGCAACCGCCTTCATAAAGATCACAGTCGTGTGGTTAATCTCGATAGGGATAGCAATCCCTGTCCCCATAAAGGGGATAGAGACGGACGTAGACAACCCTAACAACATCACATGCGTGCTGACAAAGGAGAGGTTCGGCGACTTCATGCTTTTCGGAAGTTTAGCCGCCTTCTTCACACCTCTAGCCATAATGATCGTAACCTACTTCCTAACCATACACGCACTACAGAAGAAGGCGTACCTTGTAAAGAACAAGCCACCTCAGAGGTTAACGTGGCTAACCGTGTCAACCGTATTCCAGCGTGACGAGACGCCGTGCAGTAGTCCTGAGAAGGTAGCCATGTTAGACGGCTCAAGGAAGGACAAGGCCCTGCCGAACAGTGGCGACGAGACCTTAATGAGAAGGACATCAACAATCGGCAAGAAGAGTGTGCAGACGATCTCAAACGAGCAGAGGGCAAGCAAGGTCCTAGGGATCGTGTTCTTCCTGTTCCTACTTATGTGGTGCCCTTTCTTCATCACCAACATCACCCTAGTCTTATGCGACTCGTGCAACCAGACAACCTTACAGATGTTATTAGAGATCTTCGTATGGATAGGATACGTATCGAGTGGAGTGAACCCTCTGGTCTACACGTTATTCAACAAGACGTTCAGGGACGCATTCGGCAGGTACATAACATGCAACTACAGGGCAACAAAGTCGGTCAAGACATTAAGGAAGCGTTCGAGTAAGATATACTTCAGGAACCCTATGGCCGAGAACAGTAAGTTCTTCAAGAAGCACGGGATAAGGAACGGAATAAACCCGGCGATGTACCAGTCACCGATGCGTTTGAGGAGTTCCACAATCCAGAGTAGCAGTATAATCCTATTAGACACACTGTTACTTACAGAGAACGAAGGGGACAAGACAGAGGAGCAGGTAAGCTACGTCTGA |
| gBL027 | 5-hydroxytryptamine receptor 2C | P28335-1 (canonical) | AAAACAATGGTCAACCTTAGGAACGCAGTTCACTCATTCCTGGTGCACTTAATCGGGCTATTAGTGTGGCAGTGCGACATATCCGTGAGTCCCGTAGCAGCCATCGTGACAGACATCTTCAACACATCGGACGGCGGACGTTTCAAGTTCCCTGACGGAGTACAGAACTGGCCTGCACTTTCAATCGTCATAATAATAATCATGACAATCGGCGGGAACATCTTAGTAATAATGGCCGTGAGTATGGAGAAGAAGTTACACAACGCAACGAACTACTTCCTAATGTCACTTGCGATCGCCGACATGTTAGTAGGATTACTTGTAATGCCTCTAAGTCTGTTAGCAATACTGTACGACTACGTATGGCCTCTTCCTAGGTACCTGTGCCCGGTATGGATCTCCCTAGACGTATTATTCAGCACGGCATCCATAATGCACTTATGCGCCATAAGTCTTGACAGGTACGTAGCCATCCGTAACCCTATCGAGCACTCACGTTTCAACAGTAGGACAAAGGCGATAATGAAGATAGCCATAGTATGGGCAATCTCCATAGGAGTGAGTGTCCCTATCCCTGTGATAGGACTGCGTGACGAGGAGAAGGTATTCGTGAACAACACAACGTGCGTCCTAAACGACCCTAACTTCGTGCTTATCGGCTCGTTCGTAGCATTCTTCATCCCTCTTACAATCATGGTGATCACGTACTGCCTAACAATCTACGTACTAAGGCGTCAGGCCTTAATGTTACTACACGGACACACGGAAGAGCCCCCTGGCCTGAGTCTAGACTTCCTAAAGTGCTGCAAGAGGAACACAGCCGAAGAGGAGAACTCAGCAAACCCTAACCAGGACCAGAACGCCAGGCGTAGGAAGAAGAAGGAGCGTCGTCCTAGGGGCACCATGCAGGCCATAAACAACGAGAGGAAGGCGTCAAAGGTCTTAGGCATAGTCTTCTTCGTGTTCCTAATCATGTGGTGCCCTTTCTTCATCACCAACATCTTATCAGTGCTATGCGAGAAGAGTTGCAACCAGAAGTTAATGGAGAAGCTGCTAAACGTGTTCGTCTGGATAGGGTACGTATGCTCGGGGATAAACCCTCTAGTGTACACACTATTCAACAAGATCTACAGGCGTGCCTTCTCCAACTACCTGCGTTGCAACTACAAGGTCGAGAAGAAGCCGCCCGTCCGTCAGATCCCTAGGGTCGCGGCCACAGCATTATCAGGACGTGAGCTTAACGTAAACATCTACAGGCACACGAACGAGCCCGTAATCGAGAAGGCCTCGGACAACGAGCCTGGAATCGAGATGCAGGTAGAGAACCTAGAGTTACCTGTCAACCCCAGTTCAGTAGTATCCGAGCGTATAAGTTCGGTATGA |
| gBL009 | 5-hydroxytryptamine receptor 4 | Q13639-1 (canonical) | AAAACAATGGATAAATTGGATGCTAATGTTTCTTCTGAAGAAGGTTTTGGTTCTGTTGAAAAAGTTGTTTTGTTGACTTTTTTGTCTACTGTTATTTTGATGGCTATTTTGGGTAATTTGTTGGTTATGGTTGCTGTTTGTTGGGATAGACAATTGAGAAAAATTAAAACTAATTATTTTATTGTTTCTTTGGCTTTTGCTGATTTGTTGGTTTCTGTTTTGGTTATGCCATTTGGTGCTATTGAATTGGTTCAAGATATTTGGATTTATGGTGAAGTTTTTTGTTTGGTTAGAACTTCTTTGGATGTTTTGTTGACTACTGCTTCTATTTTTCATTTGTGTTGTATTTCTTTGGATAGATATTATGCTATTTGTTGTCAACCATTGGTTTATAGAAATAAAATGACTCCATTGAGAATTGCTTTGATGTTGGGTGGTTGTTGGGTTATTCCAACTTTTATTTCTTTTTTGCCAATTATGCAAGGTTGGAATAATATTGGTATTATTGATTTGATTGAAAAAAGAAAATTTAATCAAAATTCTAATTCTACTTATTGTGTTTTTATGGTTAATAAACCATATGCTATTACTTGTTCTGTTGTTGCTTTTTATATTCCATTTTTGTTGATGGTTTTGGCTTATTATAGAATTTATGTTACTGCTAAAGAACATGCTCATCAAATTCAAATGTTGCAAAGAGCTGGTGCTTCTTCTGAATCTAGACCACAATCTGCTGATCAACATTCTACTCATAGAATGAGAACTGAAACTAAAGCTGCTAAAACTTTGTGTATTATTATGGGTTGTTTTTGTTTGTGTTGGGCTCCATTTTTTGTTACTAATATTGTTGATCCATTTATTGATTATACTGTTCCAGGTCAAGTTTGGACTGCTTTTTTGTGGTTGGGTTATATTAATTCTGGTTTGAATCCATTTTTGTATGCTTTTTTGAATAAATCTTTTAGAAGAGCTTTTTTGATTATTTTGTGTTGTGATGATGAAAGATATAGAAGACCATCTATTTTGGGTCAAACTGTTCCATGTTCTACTACTACTATTAATGGTTCTACTCATGTTTTGAGAGATGCTGTTGAATGTGGTGGTCAATGGGAATCTCAATGTCATCCACCAGCTACTTCTCCATTGGTTGCTGCTCAACCATCTGATACTTGA |
| gBL028 | 5-hydroxytryptamine receptor 5A | P47898-1 (canonical) | AAAACAATGGACTTACCCGTAAACCTTACATCCTTCAGCCTATCGACACCGTCCCCTTTAGAGACAAACCACAGCTTAGGAAAGGACGACCTGAGGCCTAGCTCGCCTCTACTGTCAGTCTTCGGAGTATTAATCTTAACGTTATTAGGCTTCTTAGTAGCGGCGACATTCGCCTGGAACTTACTTGTATTAGCGACAATATTACGTGTAAGGACATTCCACAGGGTCCCTCACAACTTAGTCGCCTCGATGGCAGTATCGGACGTGCTTGTAGCAGCATTAGTCATGCCCTTAAGTTTAGTCCACGAGCTATCAGGAAGGCGTTGGCAGCTGGGCAGGCGTTTATGCCAGCTGTGGATCGCCTGCGACGTGCTGTGCTGCACCGCGTCCATCTGGAACGTCACAGCAATAGCCTTAGACCGTTACTGGTCGATAACACGTCACATGGAGTACACGCTGAGGACCCGTAAGTGCGTCAGTAACGTCATGATAGCACTTACATGGGCGTTATCCGCAGTCATCAGTTTAGCGCCTTTACTATTCGGATGGGGAGAGACATACAGTGAGGGGAGTGAGGAGTGCCAGGTATCCAGGGAGCCCTCATACGCAGTGTTCTCCACAGTAGGCGCATTCTACTTACCTCTTTGCGTGGTCCTATTCGTCTACTGGAAGATCTACAAGGCAGCCAAGTTCAGGGTCGGATCGCGTAAGACAAACTCGGTGTCGCCCATATCTGAGGCCGTAGAGGTCAAGGACTCAGCGAAGCAGCCGCAGATGGTCTTCACCGTGAGGCACGCAACCGTGACATTCCAGCCGGAAGGCGACACATGGCGTGAGCAGAAGGAGCAGAGGGCAGCCCTTATGGTCGGAATATTAATAGGAGTATTCGTCCTGTGCTGGATACCCTTCTTCTTAACAGAGTTAATCAGCCCCCTTTGCAGCTGCGACATACCCGCCATCTGGAAGAGTATATTCCTATGGTTAGGCTACAGTAACAGCTTCTTCAACCCTCTAATCTACACGGCCTTCAACAAGAACTACAACTCCGCCTTCAAGAACTTCTTCTCAAGGCAGCACTAG |
| gBL029 | 5-hydroxytryptamine receptor 6 | P50406-1 (canonical) | AAAACAATGGTGCCAGAGCCTGGACCTACAGCGAATTCAACACCTGCCTGGGGAGCTGGACCCCCTTCCGCGCCGGGCGGATCAGGATGGGTCGCAGCAGCACTTTGCGTAGTGATAGCATTAACAGCCGCCGCAAACTCGCTACTAATCGCACTGATATGCACGCAGCCGGCACTAAGGAACACCTCGAACTTCTTCCTTGTATCCCTGTTCACCTCGGACTTAATGGTCGGGCTGGTGGTAATGCCACCCGCAATGTTAAACGCACTGTACGGGCGTTGGGTCTTAGCGAGGGGCCTATGCCTATTATGGACCGCATTCGACGTCATGTGCTGCTCAGCAAGTATCTTAAACCTGTGCCTAATCTCCCTAGACCGTTACTTATTAATATTATCGCCTTTACGTTACAAGTTACGTATGACACCTTTACGTGCATTAGCGCTGGTACTAGGGGCCTGGTCATTAGCAGCCTTAGCAAGCTTCCTACCTTTATTACTAGGATGGCACGAGCTAGGACACGCCCGTCCGCCTGTGCCTGGCCAGTGCAGGCTTCTGGCAAGCTTACCCTTCGTGCTTGTAGCAAGCGGACTGACATTCTTCCTGCCCTCAGGAGCGATCTGCTTCACCTACTGCAGGATCCTTTTAGCCGCGCGTAAGCAGGCCGTCCAGGTAGCGAGCTTAACAACAGGAATGGCCTCACAGGCAAGCGAGACATTACAGGTGCCGCGTACACCTAGGCCGGGCGTGGAGTCGGCAGACAGTAGGCGTCTTGCAACAAAGCACAGCAGGAAGGCCTTAAAGGCAAGTCTGACCCTTGGGATCTTACTGGGCATGTTCTTCGTCACGTGGCTACCCTTCTTCGTCGCAAACATAGTGCAGGCCGTGTGCGACTGCATATCCCCTGGGCTGTTCGACGTGCTGACATGGCTAGGCTACTGCAACTCAACAATGAACCCGATAATATACCCGCTGTTCATGCGTGACTTCAAGAGGGCCTTAGGAAGGTTTCTTCCTTGCCCTCGTTGCCCGCGTGAGAGGCAGGCGTCCCTGGCAAGCCCGAGCCTAAGGACAAGCCACTCAGGGCCTAGGCCTGGATTAAGCTTACAGCAGGTCTTACCTTTACCTCTTCCACCCGACAGTGACTCCGACTCGGACGCAGGATCAGGCGGCTCGTCAGGCTTACGTTTAACCGCCCAGCTTTTACTTCCGGGAGAGGCCACCCAGGACCCACCTCTTCCTACCCGTGCAGCAGCAGCAGTGAACTTCTTCAACATCGACCCGGCAGAGCCTGAGCTGAGGCCGCACCCTTTAGGAATACCTACGAACTGA |
| gBL030 | 5-hydroxytryptamine receptor 7 | P34969-1 (canonical) | AAAACAATGATGGATGTTAATTCTTCTGGTAGACCAGATTTGTATGGTCATTTGAGATCTTTTTTGTTGCCAGAAGTTGGTAGAGGTTTGCCAGATTTGTCTCCAGATGGTGGTGCTGATCCAGTTGCTGGTTCTTGGGCTCCACATTTGTTGTCTGAAGTTACTGCTTCTCCAGCTCCAACTTGGGATGCTCCACCAGATAATGCTTCTGGTTGTGGTGAACAAATTAATTATGGTAGAGTTGAAAAAGTTGTTATTGGTTCTATTTTGACTTTGATTACTTTGTTGACTATTGCTGGTAATTGTTTGGTTGTTATTTCTGTTTGTTTTGTTAAAAAATTGAGACAACCATCTAATTATTTGATTGTTTCTTTGGCTTTGGCTGATTTGTCTGTTGCTGTTGCTGTTATGCCATTTGTTTCTGTTACTGATTTGATTGGTGGTAAATGGATTTTTGGTCATTTTTTTTGTAATGTTTTTATTGCTATGGATGTTATGTGTTGTACTGCTTCTATTATGACTTTGTGTGTTATTTCTATTGATAGATATTTGGGTATTACTAGACCATTGACTTATCCAGTTAGACAAAATGGTAAATGTATGGCTAAAATGATTTTGTCTGTTTGGTTGTTGTCTGCTTCTATTACTTTGCCACCATTGTTTGGTTGGGCTCAAAATGTTAATGATGATAAAGTTTGTTTGATTTCTCAAGATTTTGGTTATACTATTTATTCTACTGCTGTTGCTTTTTATATTCCAATGTCTGTTATGTTGTTTATGTATTATCAAATTTATAAAGCTGCTAGAAAATCTGCTGCTAAACATAAATTTCCAGGTTTTCCAAGAGTTGAACCAGATTCTGTTATTGCTTTGAATGGTATTGTTAAATTGCAAAAAGAAGTTGAAGAATGTGCTAATTTGTCTAGATTGTTGAAACATGAAAGAAAAAATATTTCTATTTTTAAAAGAGAACAAAAAGCTGCTACTACTTTGGGTATTATTGTTGGTGCTTTTACTGTTTGTTGGTTGCCATTTTTTTTGTTGTCTACTGCTAGACCATTTATTTGTGGTACTTCTTGTTCTTGTATTCCATTGTGGGTTGAAAGAACTTTTTTGTGGTTGGGTTATGCTAATTCTTTGATTAATCCATTTATTTATGCTTTTTTTAATAGAGATTTGAGAACTACTTATAGATCTTTGTTGCAATGTCAATATAGAAATATTAATAGAAAATTGTCTGCTGCTGGTATGCATGAAGCTTTGAAATTGGCTGAAAGACCAGAAAGACCAGAATTTGTTTTGAGAGCTTGTACTAGAAGAGTTTTGTTGAGACCAGAAAAAAGACCACCAGTTTCTGTTTGGGTTTTGCAATCTCCAGATCATCATAATTGGTTGGCTGATAAAATGTTGACTACTGTTGAAAAAAAAGTTATGATTCATGATTAA |

**Supplementary Table S6:** Overview of primers used, including description, amplification template they were used with, new plasmid to be created, description, purpose, antibiotic and yeast marker, and reference if used previously.

| **Primer number** | **Description** | **Amplification template** | **For creation of plasmid** | **Sequence** | **Reference** |
| --- | --- | --- | --- | --- | --- |
| pBL062 | 5-HT1A fwd | 5-HT1A gBlock | cBL030 | ATCTGTCAUAAAACAATGGACGTATTATCACCTGG |  |
| pBL063 | 5-HT1A rev | 5-HT1A gBlock | cBL030 | CACGCGAUCCTCTCAATAGGGAAAACTGGCCCTACTGACGGCAGAACTTGCAC |  |
| pBL066 | 5-HT4 fwd | 5-HT4 gBlock | cBL038 | ATCTGTCAUAAAACAATGGATAAATTGGATGCTAATGTTTC |  |
| pBL067 | 5-HT4 rev | 5-HT4 gBlock | cBL038, cBL047 | CACGCGAUCCTCTCAATAGGGAAAACTGGCCTCAAGTATCAGATGGTTGAGCAGC |  |
| pBL074 | pCCW12 fwd |  | cBL030-041, cBL047 | CGTGCGAUGCTATACTGGGAGGCGTGCCAGGAACCAGGGCAAAGCAAAATAAAAG |  |
| pBL075 | pCCW12 rev |  | cBL030-041 | ATGACAGAU TATTGATATAGTGTTTAAGCGAATGACAGAAG |  |
| pBL106 | 5-HT1B fwd | 5-HT1B gBlock | cBL031 | ATCTGTCAU AAAACAATGGAAGAACCAGGAGC |  |
| pBL107 | 5-HT1B rev | 5-HT1B gBlock | cBL031 | CACGCGAU CCTCTCAATAGGGAAAACTGGCC CTAACTGGTACACTTAAACCTTATTAGTTTATG |  |
| pBL108 | 5-HT1D fwd | 5-HT1D gBlock | cBL032 | ATCTGTCAU AAAACAATGTCTCCATTGAATCAATCTG |  |
| pBL109 | 5-HT1D rev | 5-HT1D gBlock | cBL032 | CACGCGAU CCTCTCAATAGGGAAAACTGGCC TTAAGAAGCTTTTCTAAATGGAACAATTTTTTG |  |
| pBL110 | 5-HT1E fwd | 5-HT1E gBlock | cBL033 | ATCTGTCAU AAAACAATGAATATTACTAATTGTACTACTGAAGC |  |
| pBL111 | 5-HT1E rev | 5-HT1E gBlock | cBL033 | CACGCGAU CCTCTCAATAGGGAAAACTGGCC TTAAGTATGTTCTCTACATCTAATCAATTTTTTAAAAG |  |
| pBL112 | 5-HT1F fwd | 5-HT1F gBlock | cBL034 | ATCTGTCAU AAAACAATGGATTTTTTGAATTCTTCTGATC |  |
| pBL113 | 5-HT1F rev | 5-HT1F gBlock | cBL034 | CACGCGAU CCTCTCAATAGGGAAAACTGGCC TTAACATCTACATCTAACCAATTTTTGAAAAGC |  |
| pBL114 | 5-HT2A fwd | 5-HT2A gBlock | cBL035 | ATCTGTCAU AAAACAATGGACATATTATGCGAGGAG |  |
| pBL115 | 5-HT2A rev | 5-HT2A gBlock | cBL035 | CACGCGAU CCTCTCAATAGGGAAAACTGGCC TCACACGCATGATACCTTCTCG |  |
| pBL116 | 5-HT2B fwd | 5-HT2B gBlock | cBL036 | ATCTGTCAU AAAACAATGGCACTGAGTTACAGG |  |
| pBL117 | 5-HT2B rev | 5-HT2B gBlock | cBL036 | CACGCGAU CCTCTCAATAGGGAAAACTGGCC TCAGACGTAGCTTACCTGCTC |  |
| pBL118 | 5-HT2C fwd | 5-HT2C gBlock | cBL037 | ATCTGTCAU AAAACAATGGTCAACCTTAGGAAC |  |
| pBL119 | 5-HT2C rev | 5-HT2C gBlock | cBL037 | CACGCGAU CCTCTCAATAGGGAAAACTGGCC TCATACCGAACTTATACGCTCGG |  |
| pBL120 | 5-HT5A fwd | 5-HT5A gBlock | cBL039 | ATCTGTCAU AAAACAATGGACTTACCCGTAAACC |  |
| pBL121 | 5-HT5A rev | 5-HT5A gBlock | cBL039 | CACGCGAU CCTCTCAATAGGGAAAACTGGCC CTAGTGCTGCCTTGAGAAGAAG |  |
| pBL122 | 5-HT6 fwd | 5-HT6 gBlock | cBL040 | ATCTGTCAU AAAACAATGGTGCCAGAGCC |  |
| pBL123 | 5-HT6 rev | 5-HT6 gBlock | cBL040 | CACGCGAU CCTCTCAATAGGGAAAACTGGCC TCAGTTCGTAGGTATTCCTAAAGGG |  |
| pBL124 | 5-HT7 fwd | 5-HT7 gBlock | cBL041 | ATCTGTCAU AAAACAATGATGGATGTTAATTCTTCTGG |  |
| pBL125 | 5-HT7 rev | 5-HT7 gBlock | cBL041 | CACGCGAU CCTCTCAATAGGGAAAACTGGCC TTAATCATGAATCATAACTTTTTTTTCAACAGTAG |  |
| EHS0001 | HT4 (15G>R) | yBL047 | pEHS02 | AGGTTTTAGAUCTGTTGAAAAAGTTGTTTTGTTGACTTTTTTGTCTACTG |  |
| EHS0002 | HT4 (15G>R) | yBL047 | pEHS02 | ATCTAAAACCUTCTTCAGAAGAAACATTAGCATCCAATTTATCCATTGTTTT |  |
| pBL241 | HT4 (372C>Y) | yBL047 | pEHS01 | ATCTCAATATCAUCCACCAGCTACTTCTCCATTGG |  |
| pBL242 | HT4 (372C>Y) | yBL047 | pEHS01 | ATGATATTGAGAUTCCCATTGACCACCACATTCAAC |  |
| EHS0005 | HT4 (24T>M) | yBL047 | pEHS03 | AGTTGTTTTGTTGATGUTTTTGTCTACTGTTATTTTGATGGCTATTTT |  |
| EHS0006 | HT4 (24T>M) | yBL047 | pEHS03 | ACATCAACAAAACAACUTTTTCAACAGAACCAAAACCTTCTTCAGAAGAAAC |  |
| EHS0007 | HT4 (187M>T) | yBL047 | pEHS04 | ACTGTTAAUAAACCATATGCTATTACTTGTTCTGTTGTTGCTTTTT |  |
| EHS0008 | HT4 (187M>T) | yBL047 | pEHS04 | ATTAACAGUAAAAACACAATAAGTAGAATTAGAATTTTGATTAAATTTTCTTTTTTCAATCAAATCAA |  |
| EHS0009 | HT4 (260T>N) | yBL047 | pEHS05 | AAATTTGTGTAUTATTATGGGTTGTTTTTGTTTGTGTTGGGCTC |  |
| EHS0010 | HT4 (260T>N) | yBL047 | pEHS05 | ATACACAAATTUTTAGCAGCTTTAGTTTCAGTTCTCATTCTATGAGTAGA |  |
| EHS0011 | HT4 (321R>C) | yBL047 | pEHS09 | ATGTGCTTTTTUGATTATTTTGTGTTGTGATGATGAAAGATATAGAAGACCATCT |  |
| EHS0012 | HT4 (321R>C) | yBL047 | pEHS09 | AAAAAAGCACAUCTAAAAGATTTATTCAAAAAAGCATACAAAAATGGATTCAAACCAGAATTAATATAACCC |  |
| EHS0013 | HT4 (373H>P) | yBL047 | pEHS10 | ATGTCCACCACCAGCUACTTCTCCATTGGTTGCTGCTCAACC |  |
| EHS0014 | HT4 (373H>P) | yBL047 | pEHS10 | AGCTGGTGGTGGACAUTGAGATTCCCATTGACCACCACATTCA |  |
| EHS0015 | HT4 (231R>W) | yBL047 | pEHS11 | ATGGGCTGGUGCTTCTTCTGAATCTAGACCACAATCTGCT |  |
| EHS0016 | HT4 (231R>W) | yBL047 | pEHS11 | ACCAGCCCAUTGCAACATTTGAATTTGATGAGCATGTTCTTTAGC |  |
| EHS0017 | HT4 (27S>L) | yBL047 | pEHS12 | ACTTTTTTGTTGACUGTTATTTTGATGGCTATTTTGGGTAATTTGTTGGTTATGG |  |
| EHS0018 | HT4 (27S>L) | yBL047 | pEHS12 | AGTCAACAAAAAAGUCAACAAAACAACTTTTTCAACAGAACCAAAACCT |  |
| EHS0019 | HT4 (137R>H) | yBL047 | pEHS13 | ATTGCATATUGCTTTGATGTTGGGTGGTTGTTGGG |  |
| EHS0020 | HT4 (137R>H) | yBL047 | pEHS13 | AATATGCAAUGGAGTCATTTTATTTCTATAAACCAATGGTTGACAACAAAT |  |
| EHS0021 | HT4 (255T>I) | yBL047 | pEHS14 | AAATTAAAGCUGCTAAAACTTTGTGTATTATTATGGGTTGTTTTTGTTTGTG |  |
| EHS0022 | HT4 (255T>I) | yBL047 | pEHS14 | AGCTTTAATTUCAGTTCTCATTCTATGAGTAGAATGTTGATCAGCAG |  |
| EHS0025 | HT4 (302Y>F) | yBL047 | pEHS23 | TGGTTGGGTTTTATTAATTCTGGTTTGAATCCATTTTTGTATGCTTTTTTGAATAAAT |  |
| EHS0026 | HT4 (302Y>F) | yBL047 | pEHS23 | TTAATAAAACCCAACCACAAAAAAGCAGTCCAAACTTGACCTGGAACA |  |
| EHS0027 | HT4 (263I>N) | yBL047 | pEHS16 | ACTTTGTGTAATAUTATGGGTTGTTTTTGTTTGTGTTGGGCT |  |
| EHS0028 | HT4 (263I>N) | yBL047 | pEHS16 | ATATTACACAAAGUTTTAGCAGCTTTAGTTTCAGTTCTCATTCTATGAGTAGA |  |
| EHS0029 | HT4 (348T>N) | yBL047 | pEHS17 | ATGTTCTAATACUACTATTAATGGTTCTACTCATGTTTTGAGAGATGCTG |  |
| EHS0030 | HT4 (348T>N) | yBL047 | pEHS17 | AGTATTAGAACAUGGAACAGTTTGACCCAAAATAGATGGTCTTCT |  |
| EHS0031 | HT4 (137R>C) | yBL047 | pEHS18 | ATTGTGTATTGCUTTGATGTTGGGTGGTTGTTGGGTTATTCC |  |
| EHS0032 | HT4 (137R>C) | yBL047 | pEHS18 | AGCAATACACAAUGGAGTCATTTTATTTCTATAAACCAATGGTTGACAACAAAT |  |
| EHS0033 | HT4 (361A>V) | yBL047 | pEHS19 | ATGTTGTTGAATGUGGTGGTCAATGGGAATCTCAATGTCATCC |  |
| EHS0034 | HT4 (361A>V) | yBL047 | pEHS19 | ACATTCAACAACAUCTCTCAAAACATGAGTAGAACCATTAATAGTAGTAGTAGAACA |  |
| EHS0035 | HT4 (258A>T) | yBL047 | pEHS20 | AGCTACTAAAACUTTGTGTATTATTATGGGTTGTTTTTGTTTGTGTTGGG |  |
| EHS0036 | HT4 (258A>T) | yBL047 | pEHS20 | AGTTTTAGTAGCUTTAGTTTCAGTTCTCATTCTATGAGTAGAATGTTGATCAGC |  |
| EHS0037 | HT4 (214R>H) | yBL047 | pEHS21 | ATCATATTTATGTUACTGCTAAAGAACATGCTCATCAAATTCAAATGTT |  |
| EHS0038 | HT4 (214R>H) | yBL047 | pEHS21 | AACATAAATATGAUAATAAGCCAAAACCATCAACAAAAATGGAATATAAAAAGCA |  |
| EHS0039 | HT4 (223A>D) | yBL047 | pEHS22 | ATGATCATCAAATUCAAATGTTGCAAAGAGCTGGTGCTTCT |  |
| EHS0040 | HT4 (223A>D) | yBL047 | pEHS22 | AATTTGATGATCAUGTTCTTTAGCAGTAACATAAATTCTATAATAAGCCAAAACCATCA |  |
| ID350 | RnPTS<-TEF1-PGK1->RnSPR cassette | pCfB1251 | pDAM23 | CGTGCGAUTTATTCTCCTTTGTAGACCACAAT | (Germann et al., 2016) |
| ID389 | RnPTS<-TEF1-PGK1->RnSPR cassette | pCfB1251 | pDAM23 | CACGCGAUTTAAATGTCATAGAAGTCCACGTG | (Germann et al., 2016) |
| ID2149 | PaPCBD1<-TEF1-PGK1->RatDHPR | pCfB1248 | pDAM24 | CGTGCGAUTTACTTTCTACCTTCAGCAG | (Germann et al., 2016) |
| ID2153 | PaPCBD1<-TEF1-PGK1->RatDHPR | pCfB1248 | pDAM24 | CACGCGAUTTAGAAGTAAGCTGGAGTC | (Germann et al., 2016) |
| TJOS-62 | gRNA targeting X-4 site | pCfB3042 | pDAM22 | CGTGCGAUagggaacaaaagctggagct | (Jakočiūnas et al., 2015) |
| TJOS-63 | gRNA targeting X-4 site | pCfB3042 | pDAM22 | AGTGCAGGUagggaacaaaagctggagct | (Jakočiūnas et al., 2015) |
| TJOS-64 | gRNA targeting XI-3 site | pCfB3045 | pDAM22 | ATCTGTCAUagggaacaaaagctggagct | (Jakočiūnas et al., 2015) |
| TJOS-65 | gRNA targeting XI-3 site | pCfB3045 | pDAM22 | CACGCGAUtaactaattacatgactcga | (Jakočiūnas et al., 2015) |
| TJOS-66 | gRNA targeting XII-4 site | pCfB3049 | pDAM22 | ACCTGCACUtaactaattacatgactcga | (Jakočiūnas et al., 2015) |
| TJOS-67 | gRNA targeting XII-4 site | pCfB3049 | pDAM22 | ATGACAGAUtaactaattacatgactcga | (Jakočiūnas et al., 2015) |

**Supplementary Table S7: Overview of plasmids used, including plasmid name, description, purpose, antibiotic and yeast marker, and reference if used previously.**

| **Plasmid name** | **Description** | **Purpose** | **Antibiotic marker** | **Yeast marker** | **Reference** |
| --- | --- | --- | --- | --- | --- |
| cBL030 | tADH1->CCW12->HT1A(Sc)->tCyC1 | expression of 5-HTx | AMP | HIS | / |
| cBL031 | tADH1->CCW12->HT1B(Hs)->tCyC1 | expression of 5-HTx | AMP | HIS | / |
| cBL032 | tADH1->CCW12->HT1D(Sc)->tCyC1 | expression of 5-HTx | AMP | HIS | / |
| cBL033 | tADH1->CCW12->HT1E(Sc)->tCyC1 | expression of 5-HTx | AMP | HIS | / |
| cBL034 | tADH1->CCW12->HT1F(Sc)->tCyC1 | expression of 5-HTx | AMP | HIS | / |
| cBL035 | tADH1->CCW12->HT2A(Sc)->tCyC1 | expression of 5-HTx | AMP | HIS | / |
| cBL036 | tADH1->CCW12->HT2B(Sc)->tCyC1 | expression of 5-HTx | AMP | HIS | / |
| cBL037 | tADH1->CCW12->HT2C(Sc)->tCyC1 | expression of 5-HTx | AMP | HIS | / |
| cBL038 | tADH1->CCW12->HT4(Sc)->tCyC1 | expression of 5-HTx | AMP | HIS | / |
| cBL039 | tADH1->CCW12->HT5(Sc)->tCyC1 | expression of 5-HTx | AMP | HIS | / |
| cBL040 | tADH1->CCW12->HT6(Sc)->tCyC1 | expression of 5-HTx | AMP | HIS | / |
| cBL041 | tADH1->CCW12->HT7(Sc)->tCyC1 | expression of 5-HTx | AMP | HIS | / |
| cBL047 | pCCW-HT4 for integration into XII-4 | integration of 5-HT4 | AMP | none | / |
| pCfB6908 | gRNA plasmid for XII-4 | gRNA | AMP | LEU | / |
| pTAJAK161 | Cas9 plasmid | Cas9 | AMP | HIS | / |
| pCfB9221 | HsDDC<-PTDH3-PTEF1->SmTPH for XI-3 integration | for integration | AMP | none | (Germann et al., 2016) |
| pDAM23 | RnPTS<-TEF1-PGK1->RnSPR for X-4 integration | for integration | AMP | none | / |
| pDAM24 | PaPCBD1<-TEF1-PG‚K1->RnDHPR for XII-4 integration | for integration | AMP | none | / |
| pDAM22 | XI-3, X-4, XII-4 gRNA plasmid | gRNA plasmid | CloNAT | none | / |
| pCfB2772 | pTY2-KlURA3-TAG-PGK1-_SmTPH for TY2 integration | for TY2 integration | AMP | none | (Germann et al., 2016) |
| pEHS01 | tADH1->CCW12->HT4 372C>Y ->tCYC1 | for integration | AMP | none | / |
| pEHS02 | tADH1->CCW12->HT4 15G>R->tCYC1 | for integration | AMP | none | / |
| pEHS03 | tADH1->CCW12->HT4 24T>M->tCYC1 | for integration | AMP | none | / |
| pEHS04 | tADH1->CCW12->HT4 187M>T->tCYC1 | for integration | AMP | none | / |
| pEHS05 | tADH1->CCW12->HT4 260T>N->tCYC1 | for integration | AMP | none | / |
| pEHS09 | tADH1->CCW12->HT4 321R>C ->tCYC1 | for integration | AMP | none | / |
| pEHS10 | tADH1->CCW12->HT4 373H>P ->tCYC1 | for integration | AMP | none | / |
| pEHS11 | tADH1->CCW12->HT4 231R>W ->tCYC1 | for integration | AMP | none | / |
| pEHS12 | tADH1->CCW12->HT4 27S>L ->tCYC1 | for integration | AMP | none | / |
| pEHS13 | tADH1->CCW12->HT4 137R>H ->tCYC1 | for integration | AMP | none | / |
| pEHS14 | tADH1->CCW12->HT4 255T>I ->tCYC1 | for integration | AMP | none | / |
| pEHS15 | tADH1->CCW12->HT4 364C>R ->tCYC1 | for integration | AMP | none | / |
| pEHS16 | tADH1->CCW12->HT4 263I>N ->tCYC1 | for integration | AMP | none | / |
| pEHS17 | tADH1->CCW12->HT4 348T>N ->tCYC1 | for integration | AMP | none | / |
| pEHS18 | tADH1->CCW12->HT4 137R>C ->tCYC1 | for integration | AMP | none | / |
| pEHS19 | tADH1->CCW12->HT4 361A>V ->tCYC1 | for integration | AMP | none | / |
| pEHS20 | tADH1->CCW12->HT4 258A>T ->tCYC1 | for integration | AMP | none | / |
| pEHS21 | tADH1->CCW12->HT4 214R>H ->tCYC1 | for integration | AMP | none | / |
| pEHS22 | tADH1->CCW12->HT4 223A>D ->tCYC1 | for integration | AMP | none | / |
| pEHS23 | tADH1->CCW12->HT4 302Y>F -> tCYC1 | for integration | AMP | none | / |
| pCfB1248 | pX-4-LoxP-SpHIS5-PaPCBD1<-TEF1-PGK1->RatDHPR | template for pDAM24 | AMP | HIS | (Germann et al., 2016) |
| pCfB1251 | pX-3-LoxP-KlLEU2-RnPTS<-TEF1-PGK1->RnSPR | template for pDAM23 | AMP | LEU | (Germann et al., 2016) |
| pCfB3040 | USER plasmid for XII-4 integration | vector for 5-HT4 integration plasmids | AMP | none | (Jessop-Fabre et al., 2016) |
| pCfB3042 | gRNA targeting X-4 site | template for gRNA plasmid pDAM22 | AMP | none | (Jessop-Fabre et al., 2016) |
| pCfB3045 | gRNA targeting XI-3 site | template for gRNA plasmid pDAM22 | AMP | none | (Jessop-Fabre et al., 2016) |
| pCfB3049 | gRNA targeting XII-4 site | template for gRNA plasmid pDAM22 | AMP | none | (Jessop-Fabre et al., 2016) |

**Supplementary Table S8:** Overview of yeast strains used, including strain name, description, plasmid used for integration, in which Figure it was used, and reference if used previously.

| **Strain Name** | **Description** | **Plasmid integrated** | **Precursor strain** | **Figure** | **Reference** |
| --- | --- | --- | --- | --- | --- |
| yWS677 | sst2Δ0 far1Δ0 bar1Δ0 ste2Δ0 ste12Δ0 gpa1Δ0 ste3Δ0 mf(alpha)1Δ0 mf(alpha)2Δ0 mfa1Δ0 mfa2Δ0 gpr1Δ0 gpa2Δ0 | / | BY4741 | not used in study, for genetic information only | (Shaw et al., 2019) |
| yWS2261 | Design 4 without receptor - GPA1 | / | yWS677 | Base strain | (Shaw et al., 2019) |
| yWS2262 | Design 4 without receptor - Gαs/olf | / | yWS677 | Base strain | (Shaw et al., 2019) |
| yWS2263 | Design 4 without receptor - Gα12 | / | yWS677 | Base strain | (Shaw et al., 2019) |
| yWS2264 | Design 4 without receptor - Gα13 | / | yWS677 | Base strain | (Shaw et al., 2019) |
| yWS2265 | Design 4 without receptor - Gαi1/2 | / | yWS677 | Base strain | (Shaw et al., 2019) |
| yWS2266 | Design 4 without receptor - Gαi3 | / | yWS677 | Base strain | (Shaw et al., 2019) |
| yWS2267 | Design 4 without receptor - Gαz | / | yWS677 | Base strain | (Shaw et al., 2019) |
| yWS2268 | Design 4 without receptor - Gα15/16 | / | yWS677 | Base strain | (Shaw et al., 2019) |
| yWS2269 | Design 4 without receptor - Gαq/11 | / | yWS677 | Base strain | (Shaw et al., 2019) |
| yWS2270 | Design 4 without receptor - Gα14 | / | yWS677 | Base strain | (Shaw et al., 2019) |
| yWS2271 | Design 4 without receptor - Gαo | / | yWS677 | Base strain | (Shaw et al., 2019) |
| yWS2272 | Design 4 without receptor - tGPA1 | / | yWS677 | Base strain | (Shaw et al., 2019) |
| 144x 5-HT4 library | 12x HT receptor in 12 G alpha backgrounds |  | yWS2261-yWS2272 | Fig. 1C | / |
| yBL161 | 5-HT4 in XII-4 - GPA1 | yBL030 | yWS2261 | Fig. 2 | / |
| yBL162 | 5-HT4 in XII-4 - Gαs/olf | yBL031 | yWS2262 | Fig. 2 | / |
| yBL163 | 5-HT4 in XII-4 - Gα12 | yBL032 | yWS2263 | Fig. 2 | / |
| yBL164 | 5-HT4 in XII-4 - Gα13 | yBL033 | yWS2264 | Fig. 2 | / |
| yBL165 | 5-HT4 in XII-4 - Gαi1/2 | yBL034 | yWS2265 | Fig. 2 | / |
| yBL166 | 5-HT4 in XII-4 - Gαi3 | yBL035 | yWS2266 | Fig. 2 | / |
| yBL167 | 5-HT4 in XII-4 - Gαz | yBL036 | yWS2267 | Fig. 2, Fig. 3B, 3C, 3D, 3F | / |
| yBL168 | 5-HT4 in XII-4 - Gα15/16 | yBL037 | yWS2268 | Fig. 2 | / |
| yBL169 | 5-HT4 in XII-4 - Gαq/11 | yBL038 | yWS2269 | Fig. 2 | / |
| yBL170 | 5-HT4 in XII-4 - Gα14 | yBL039 | yWS2270 | Fig. 2 | / |
| yBL171 | 5-HT4 in XII-4 - Gαo | yBL040 | yWS2271 | Fig. 2 | / |
| yBL172 | 5-HT4 in XII-4 - tGPA1 | yBL041 | yWS2272 | Fig. 2 | / |
| yBL241 | 5-HT4 (372C>Y) in XII-4 - GPA1 | pEHS01 | yWS2261 | Fig. 4 | / |
| yBL242 | 5-HT4 (15G>R) in XII-4 - GPA1 | pEHS02 | yWS2261 | Fig. 4 | / |
| yBL243 | 5-HT4 (24T>M) in XII-4 - GPA1 | pEHS03 | yWS2261 | Fig. 4 | / |
| yBL244 | 5-HT4 (187M>T) in XII-4 - GPA1 | pEHS04 | yWS2261 | Fig. 4 | / |
| yBL245 | 5-HT4 (260T>N) in XII-4 - GPA1 | pEHS05 | yWS2261 | Fig. 4 | / |
| yBL246 | 5-HT4 (321R>C) in XII-4 - GPA1 | pEHS09 | yWS2261 | Fig. 4 | / |
| yBL247 | 5-HT4 (373H>P) in XII-4 - GPA1 | pEHS10 | yWS2261 | Fig. 4 | / |
| yBL248 | 5-HT4 (231R>W) in XII-4 - GPA1 | pEHS11 | yWS2261 | Fig. 4 | / |
| yBL249 | 5-HT4 (27S>L) in XII-4 - GPA1 | pEHS12 | yWS2261 | Fig. 4 | / |
| yBL250 | 5-HT4 (137R>H) in XII-4 - GPA1 | pEHS13 | yWS2261 | Fig. 4 | / |
| yBL251 | 5-HT4 (255T>I) in XII-4 - GPA1 | pEHS14 | yWS2261 | Fig. 4 | / |
| yBL253 | 5-HT4 (263I>N ) in XII-4 - GPA1 | pEHS16 | yWS2261 | Fig. 4 | / |
| yBL254 | 5-HT4 (348T>N) in XII-4 - GPA1 | pEHS17 | yWS2261 | Fig. 4 | / |
| yBL255 | 5-HT4 (137R>C) in XII-4 - GPA1 | pEHS18 | yWS2261 | Fig. 4 | / |
| yBL256 | 5-HT4 (361A>V) in XII-4 - GPA1 | pEHS19 | yWS2261 | Fig. 4 | / |
| yBL257 | 5-HT4 (258A>T ) in XII-4 - GPA1 | pEHS20 | yWS2261 | Fig. 4 | / |
| yBL258 | 5-HT4 (214R>H) in XII-4 - GPA1 | pEHS21 | yWS2261 | Fig. 4 | / |
| yBL259 | 5-HT4 (223A>D ) in XII-4 - GPA1 | pEHS22 | yWS2261 | Fig. 4 | / |
| yBL260 | 5-HT4 (302Y>F) in XII-4 - GPA1 | pEHS23 | yWS2261 | Fig. 4 | / |
